# Short-term responses to ocean acidification: effects on relative abundance of eukaryotic plankton from the tropical Timor Sea

**DOI:** 10.1101/2020.04.30.068601

**Authors:** Janina Rahlff, Sahar Khodami, Lisa Voskuhl, Matthew P. Humphreys, Christian Stolle, Pedro Martinez Arbizu, Oliver Wurl, Mariana Ribas-Ribas

**Author notes:** University of Duisburg-Essen, Department of Chemistry, Environmental Microbiology and Biotechnology (EMB), 45141 Essen, Germany. Project management Jülich, 18069 Rostock, Germany (Christian Stolle).

## Abstract

Anthropogenic carbon dioxide (CO_2_) emissions drive climate change and pose one of the major challenges of our century. The effects of increased CO_2_ in the form of ocean acidification (OA) on the communities of marine planktonic eukaryotes in tropical regions such as the Timor Sea are barely understood. Here, we show the effects of high CO_2_ (*p*CO_2_=1823±161 μatm, pH_T_=7.46±0.05) versus *in situ* CO_2_ (*p*CO_2_=504±42 μatm, pH_T_=7.95±0.04) seawater on the community composition of marine planktonic eukaryotes immediately and after 48 hours of treatment exposure in a shipboard microcosm experiment. Illumina sequencing of the V9 hypervariable region of 18S rRNA (gene) was used to study the eukaryotic community composition. Down-regulation of extracellular carbonic anhydrase occurred faster in the high CO_2_ treatment. Increased CO_2_ significantly suppressed the relative abundances of different eukaryotic operational taxonomic units (OTUs), including important primary producers. These effects were consistent between abundant (DNA-based) and active (cDNA-based) taxa after 48 hours, e.g., for the diatoms *Trieres chinensis* and *Stephanopyxis turris*. Effects were also very species-specific among different diatoms. Planktonic eukaryotes showed adaptation to the CO_2_ treatment over time, but many OTUs were adversely affected by decreasing pH. OA effects might fundamentally impact the base of marine biodiversity, suggesting profound outcomes for food web functioning in the future ocean.

## 1. INTRODUCTION

The world’s oceans are a major sink for anthropogenic carbon dioxide (CO_2_) (Friedlingstein et al. 2019, Gruber et al. 2019). Absorption of atmospheric CO_2_, which is currently at 414 μatm (February 2020, https://www.co2.earth/) and driven by ongoing anthropogenic emissions, decreases pH while increasing the bicarbonate ion concentration and partial pressure of CO_2_ (*p*CO_2_) of seawater. Man-made CO_2_ emissions are expected to lead to ocean pH changes greater than any experienced in the last 300 million years, with a maximum predicted pH reduction of 0.77 in surface water for the year 2300 (Caldeira & Wickett 2003). The decrease in seawater pH, known as ocean acidification (OA) or “the other CO_2_ problem” (Doney et al. 2009), affects ocean biogeochemistry and the physiology of a plethora of marine organisms, especially the calcifying ones (Orr et al. 2005, Kaniewska et al. 2012, Schlüter et al. 2016). The fifth assessment report of the IPCC suggests that net primary productivity in the tropical ocean will most likely decline by 7-16 % by 2081-2100 assuming the worst case “business as usual” scenario (Representative Concentration Pathway=RCP8.5), which predicts an atmospheric CO_2_ concentration of >1000 μatm by the end of this century (Pachauri et al. 2014). Moreover, profound implications of OA effects on community and ecosystem processes can be expected but are difficult to predict (Riebesell et al. 2007, Fabry et al. 2008).

Changes in seawater pH and *p*CO_2_ may have critical effects on the primary producers, i.e., phytoplankton that convert CO_2_ to organic carbon during photosynthesis, and thus affect the base of the marine food web. Increased CO_2_ is known to promote phytoplankton growth and photosynthetic rates (Wolf-Gladrow et al. 1999, Kroeker et al. 2013), although increases in the latter are assumed to be small and dependent on the phytoplankton species under investigation (reviewed by Raven et al. (2005)). Typical acclimation effects of phytoplankton to elevated CO_2_ include the down-regulation of carbon concentrating mechanisms, e.g., of the enzyme carbonic anhydrase (Mustaffa et al. 2017, Deppeler et al. 2018), demonstrating direct effects of OA on intracellular pH and enzyme activities. Furthermore, CO_2_-driven changes in element stoichiometry (King et al. 2015), community structure (Tortell et al. 2002, Feng et al. 2010), and size classes (Suffrian et al. 2008) of primary producers can alter food availability and quality for grazers and cascade up the whole food web (Rossoll et al. 2012, Cripps et al. 2016).

Copepods (phylum Arthropoda, class Copepoda (Pancrustacea)) are the most abundant metazoans on Earth (Humes 1994). They form the major dietary link between phytoplankton and ichthyoplankton as well as larger fish (Turner 2004). Copepods and other crustaceans were assumed to be more resilient towards OA effects compared to other marine organisms (Kurihara & Ishimatsu 2008, Whiteley 2011). However, this assumption mainly results from studies on adult copepod females and has been questioned (Cripps et al. 2014) supporting the lack of conclusive knowledge about OA effects on copepods that exist in the literature.

Insights on the effects of lowered pH on natural marine nano- to meso-sized planktonic organisms from the tropics are scarce. Hence, our objective was to study the impact of OA on a natural plankton community from the Timor Sea, tropical Indian Ocean in a microcosm experiment that was conducted onboard the *R/V* Falkor in October 2016. Probable OA effects on planktonic communities were investigated in one treatment with reduced pH (*p*CO_2_=1823±161 μatm, pH_T_=7.46±0.05) compared to an unaltered control (*p*CO_2_=504±42 μatm, pH_T_=7.95±0.04) at two consecutive time points. The effects of OA on abundant and active planktonic organisms were investigated as inferred from DNA and cDNA-based V9 hypervariable region of the 18S rRNA (gene) sequencing, respectively. Since literature on the topic reports inconclusive results of OA effects sometimes even for the same organism (Ridgwell et al. 2009, Meyer & Riebesell 2015, Thor & Oliva 2015), we expected to see significant short-term OA effects after different exposure times on the relative abundance of some taxa as well as missing responses in others.

## 2. MATERIALS AND METHODS

### 2.1 Sampling and microcosm setup

During a research cruise on *R/V* Falkor (FK161010), sample water containing planktonic organisms was collected from the chlorophyll maximum at 65 m depth using a Conductivity, Temperature, Depth (CTD) unit at 19:20, 20^th^ October 2016 (UTC) from the Timor Sea (11°51.25’S, 127°15.23’E). The region is tropical with summer and winter sea surface temperature >26 and >22 °C, respectively, and experiences periodic cyclone-generated storm currents (James et al. 2004). Water temperature and salinity at the time of sampling were at 28 °C and 35, respectively. Sampled seawater was randomly filled from CTD bottles into 24 x 1-L Nalgene polycarbonate bottles (Thermo Fisher Scientific, Darmstadt, Germany), which were wrapped into a foil (No 298, Lee Filters, Andover, UK) filtering daylight to approach light conditions at the sampling depth. Half of the bottles were acidified using an equimolar addition of strong acid (1 M HCl) and HCO_3_ (1 M NaHCO_3_) to simulate effects of low pH/high CO_2_ (treatment HICO) according to conditions expected in a future ocean (Pachauri et al. 2014). The other half of samples was kept at *in situ* conditions (treatment ISCO). All bottles were placed into a baby pool (99 x 99 x 23 cm, Wehnke, Germany). During sample incubations, fresh oceanic water was constantly pumped into the pool to keep the temperature as similar as possible to *in situ* conditions. Pool temperature was monitored by a portable PCD650 (EuTech Instruments, Singapore, Supplementary Figure S1). Incubations were carried out from the 20^th^ (t_0_) to the 23^th^ October, 00:40 UTC (t_2_), and for each day beginning at t_0_, four replicate bottles of each the HICO and the ISCO treatment were processed. The time period from acidification until water filtration for t_0_ samples was three hours. Before water sample processing, the pH was measured in each bottle to monitor that the manipulation was effective.

### 2.2 Chlorophyll *a* extraction

Discrete chlorophyll *a* (chl *a*) analysis was performed on sampling water (initial samples after CTD, HICO, ISCO, n=4), using the fluorometer JENWAY 6285 (Bibby Scientific Ltd., Felsted, Essex, UK). Readings on standards were taken by using commercial pure chl *a* (Sigma Aldrich, Taufkirchen, Germany). Water samples (160–300 mL of the 1L) were filtered onboard the *R/V* immediately onto glass microfiber filters (GF/F, diameter: 25 mm, Whatman, UK), and were stored at −20 °C, and shipped on dry ice for further analysis in the home laboratory. The filtered samples were extracted in 90% ethanol for 24 hours in the dark, before being measured fluorometrically in triplicates and according to the EPA Method 445.0 (Arar & Collins 1997).

### 2.3 Analysis of carbonate chemistry

Dissolved inorganic carbon (DIC) and total alkalinity (TA) samples were analyzed at the University of Southampton using a VINDTA 3C (Marianda, Kiel, Germany). The DIC was measured by coulometric titration, and TA by potentiometric titration and calculated using a modified Gran plot approach as implemented by Calkulate (version 1.0.2) (Bradshaw et al. 1981, Humphreys 2015). Measurements were calibrated using certified reference material (batches 144, 151 and 160) obtained from A.G. Dickson (Scripps Institution of Oceanography, USA). The 1*σ* measurement precision was ±3 and ±2 μmol kg^−1^ for DIC and TA, respectively. The remaining carbonate chemistry variables were calculated with PyCO_2_SYS (version 1.3.0) (Lewis & Wallace 1998, Van Heuven et al. 2011, Humphreys et al. 2020) using the carbonic acid dissociation constants of Lueker et al. (2000).

The dissociation constant for bisulphate dissociation was chosen according to Dickson (1990), and the ratio of total borate to salinity was in accordance with Lee et al. (2010).

Discrete samples for inorganic nutrients (nitrate and nitrite, phosphate and silicate) were collected from the same intervals as the DIC and TA samples. Inorganic nutrients were measured using a continuous flow analyzer (Quattro, CFA), Reader, Easychem (Strickland 1968, Fanning & Pilson 1973). The detection limit was 0.4 μM for nitrate and nitrite, 0.04 μM for phosphate and 0.1 μM for silicate, and the limit of determination was 1.0 μM, 0.13 μM, and 0.3 μM respectively.

### 2.4 DNA extraction and cDNA synthesis

For DNA and RNA extraction, a two-step filtration of sampling water (150-400 mL from each bottle) was conducted, namely water was filtered through a 3.0 μm pore size membrane filter. The flow-through was further filtered onto a 0.2 μm polycarbonate membrane filter (47 mm diameter, Whatman, Maidstone, UK). All filters were initially stored at −80°C prior analysis and shipped on dry ice to the home laboratory. Extraction of DNA and RNA from t_0_ and t_2_ filters for three biological replicates (half of a 0.2 and a 3 μm filter were pooled in each case) was performed by using the DNA + RNA + Protein Extraction Kit (Roboklon, Berlin, Germany) and a modified protocol as described in Rahlff et al. (2017). Remaining DNA in RNA samples was digested on-column using 3 U of DNase, and all samples were subsequently checked for genomic DNA contaminations by polymerase chain reaction (PCR). A quantity of 10 ng RNA was reversely transcribed to cDNA using the NG dART Kit and therein included random hexamer primers (Roboklon, Berlin, Germany). Negative controls without reverse transcriptase and without RNA were included. The reaction was incubated for 60 minutes at 50°C followed by 5 minutes at 85°C.

### 2.5 Library preparation from V9-18S rRNA (gene) amplicons, sequencing and bioinformatics

Two independent PCRs were carried out to attach the Nextera XT compatible Illumina adapter overhangs at the 5’ ends of each amplicon following the Illumina 16S Metagenomic Sequencing Library Preparation guide (15044223Rev.B). The first PCR was conducted on a 3PrimeG thermocycler (Techne, Staffordshire, UK) using Phusion High Fidelity DNA Polymerase (Thermo Fisher Scientific, Darmstadt, Germany), 10 mM dNTPs, 3% DMSO and 0.5 μM of the Tara Oceans eukaryote primer set 1389F/1510R targeting the V9 hypervariable region of the 18S ribosomal RNA gene (Amaral-Zettler et al. 2009). The primers were tagged with part of the Nextera compatible Illumina adapter overhang at the 5’ ends of each primer. The PCR-cycling program was modified from a previously described one for this primer set (Alberti et al. 2017), namely 27 and 30 cycles instead of 25 were chosen for DNA and cDNA template, respectively.

Five μL of the first PCR products have been purified using 2 μL of ExoSAP-IT PCR Product Cleanup Reagent (Thermo Fisher Scientific) following the manufacturer’s protocol. A second PCR using Nextera XT V2 Indexed primers (dual indexing approach) has been performed to bind the Illumina overhang adapters to the product of the first PCR and was conducted at 98 °C for 2 min, 7 cycles of 15 s at 98 °C, 30 s at 62°C, 30 s at 72 °C and 72 °C for 2 min. The PCR products’ cleanup was performed using 60% of the PCR products volume of Agencourt^®^ Ampure^®^ XP magnetic beads (Beckmann Coulter, Brea, CA, USA), and concentrations were measured using Qubit Fluorometric Quantitation (Thermo Fisher Scientific, Darmstadt, Germany). All amplicon concentrations were subsequently adjusted to 8 nM, pooled and purified from a 2% agarose gel stained with 2% Gel Red using Monarch^®^ Nucleic Acid Purification kit (New England Biolabs, Ipswich, MA, USA). The library was sequenced in two independent runs on an Illumina MiSeq platform using the MiSeq Reagent Nano Kit v2 (150 paired-end cycles). The resulted reads have been demultiplexed by MiSeq considering Nextera index sequences for both forward and reverse strands.

BBMap tool (Bushnell 2018) has been used for trimming the primers. The Vsearch pipeline (Rognes et al. 2016) was used for making contigs from each sample using the following settings: expected mean size of 170 bp with +/- 30 bp intervals, minimum contig overlap at 25 bp, and maximum allowed differences in the contig at 15 bp. Contigs have been filtered with maximum expected errors of 0.5 and maximum of 0 ambiguities in the contig and de-replicated. All de-noised and de-replicated samples from libraries have been initially clustered at 98% similarity to produce the initial operational taxonomic units (OTUs). After excluding chimeras, the initial OTUs have been clustered at a similarity threshold of 97 % (species level) for each sample, which has been recently recommended for PCR-based high-throughput sequencing data to obtain more realistic richness and Shannon diversity data from 18S amplicon data (Wylezich et al. 2018). The generated OTUs have been blasted against GenBank to assign taxonomies. Sequence reads were deposited at NCBI’s sequence read archive (SRA) under Bioproject ID PRJNA623264.

### 2.6 Data analysis

A total number of 2435 OTUs resulted after implementing all filtrations, in which the number of unique reads per library ranged from 20616 to 41687. After de-noising, chimera detection and de-replicating this has been reduced to 3051 to 5673. Number of resulted OTUs were between 503 and 1136 for the different samples (Supplementary Table S1).

Sequence data were analyzed using the “phyloseq” package (McMurdie & Holmes 2013) in R version 3.6.1. (Team 2017). Reads were normalized to minimum sequencing depth, which corresponded to 18056 reads. One replicate of the HICO t_0_ cDNA samples was removed from the dataset due to a too small sequencing depth of 334.

For all calculations and plots we focused on the following phylogenetic groups: apicomplexans, brown-algae, cercozoans, choanoflagellates, ciliates, crustaceans, cryptomonads, ctenophores, diatoms, dinoflagellates, euglenoids, eukaryotes (no further classification), forams, golden_algae, green_algae, haptophytes, kinetoplastids and pelagophytes. Relative abundances were calculated and used for the following analyses. Shannon-Wiener diversity indices (α- diversity) were calculated using the estimate_richness function of the “phyloseq” package. We visualized the differences (β-diversity) of the eukaryotic planktonic community composition of samples through a Nonmetric Multidimensional Scaling (NMDS) plot based on Bray–Curtis dissimilarity indices.

For finding significant differences for extracellular carbonic anhydrase (eCA) concentrations and Shannon-Wiener indices between treatments, a one-way analysis of variances (one-way ANOVA) with 95 % confidence level was performed using R. The prerequisites for parametric tests, i.e., normal distribution of data and variance homogeneity were previously confirmed using Shapiro and Bartlett tests in R, respectively. A Tukey HSD *post-hoc* test was used to make multiple comparisons of means to find significant differences.

A two-sided t-test assuming unequal variances (Welch t-test) was conducted to find significant differences (α≤0.05) between relative abundances of individual OTUs in the ISCO and HICO treatments (n=3). Since the sample HICO t_0_ cDNA lacked one replicate, the comparison ISCO/HICO for t0 cDNA was omitted.

To elucidate the abundance of significantly CO_2_-affected taxa in both treatments., the proportion of significantly CO_2_-affected OTUs was calculated and assigned to one of five abundance categories (Table 2).

## 3. RESULTS

### 3.1 Carbonate chemistry and temperature

Temperature in the pool varied between 27.0 and 34.0 °C from t_0_ to t_2_ (Supplementary Figure S1). Seawater pH_T_, i.e., pH on the Total scale, was consistent at t_0_ and t_1_ under both *p*CO_2_ conditions with average values (mean ± standard deviation) of 7.44±0.03 and 7.93±0.02 for the HICO and ISCO treatment, respectively (Figure 1A). We measured a small increase in pH_T_ of circa 0.06 in both experiments at t_2_, to 7.50±0.06 and 7.99±0.03. Dissolved inorganic carbon (DIC) was relatively stable throughout all experiments under both *p*CO_2_ conditions (Figure 1B), with an average of 2018±14 μmol kg^−1^ and 2213±14 μmol kg^−1^ in the ISCO and HICO treatment, respectively.

**Figure 1:**
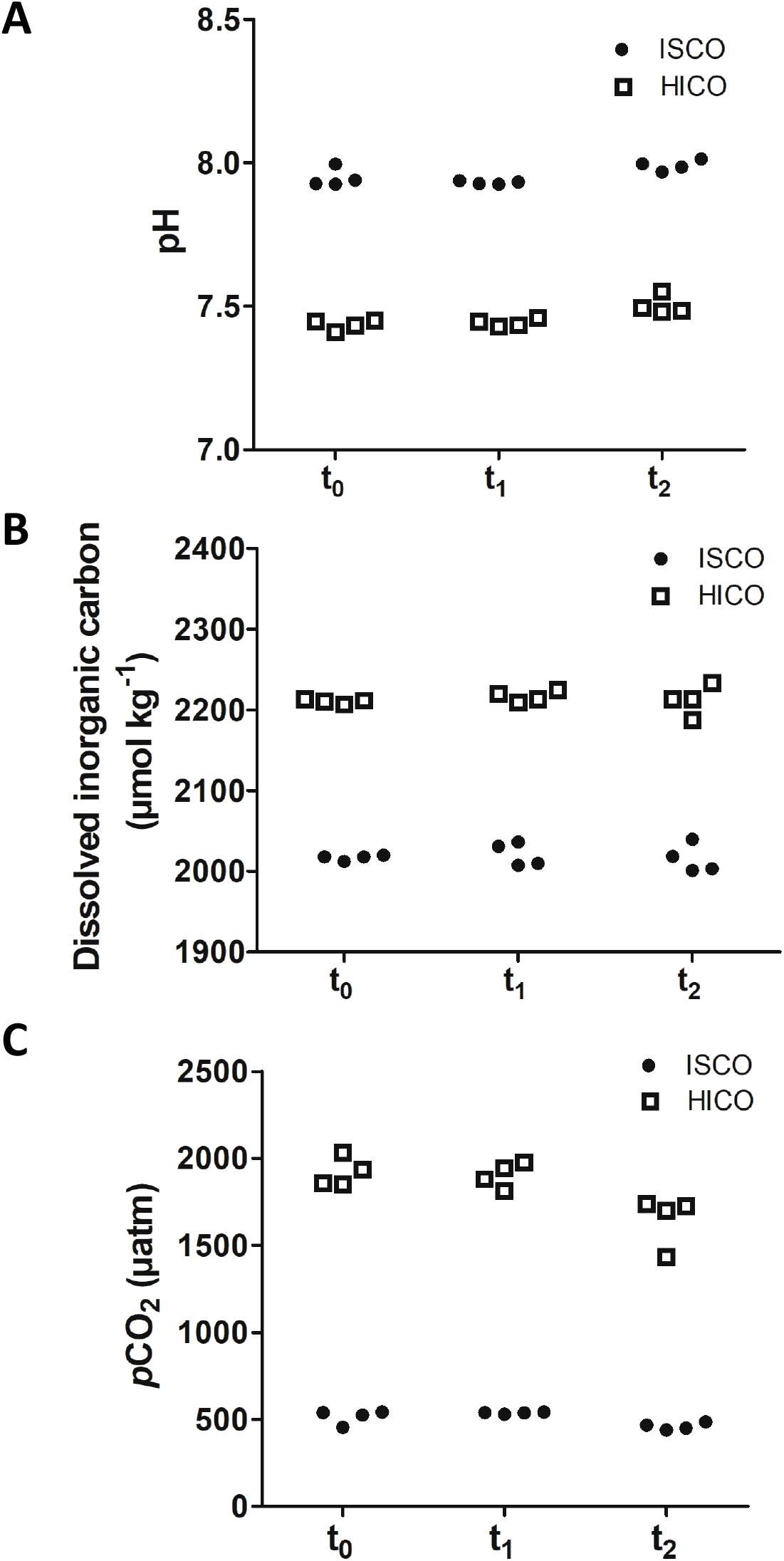
Carbonate chemistry measurements during the experiments. A) Seawater pH (Total scale) measured directly during the experiments at *in situ* temperature (c. 25 °C). B) Dissolved inorganic carbon (DIC) measured later from subsamples of the experiment media. C) Seawater *p*CO_2_ at *in situ* temperature calculated from the pH and DIC measurements shown in A) and B). For samples with no DIC measurement, the average DIC across all timepoints for all experiments at a similar CO_2_ level was used instead (i.e. 2018 μmol kg^−1^ for ISCO experiments, and 2213 μmol kg^−1^ for HICO experiments).

The seawater *p*CO_2_ was calculated from measured pH_T_ and DIC along with relevant metadata, using the mean DIC values for each CO_2_ treatment where those measurements were missing (Figure 1C). Like pH_T_, seawater *p*CO_2_ was consistent at t_0_ and t_1_ under both *p*CO_2_ conditions with average values of 1910±73 μatm and 526±30 μatm for the HICO and ISCO treatments respectively, but these values decreased to 1648±144 μatm and 460±20 μatm at t_2_. The calculated decrease in *p*CO_2_ at t_2_ was driven by the measured increase in pH_T_, which could be due to organic alkalinity degradation during the experiments (Martín Hernández-Ayon et al. 2007).

### 3.2 Chlorophyll *a*, nutrients, extracellular carbonic anhydrase

Discrete chl *a* analyses revealed concentrations <0.4 μg L^−1^ (n=4) with no consistent difference between HICO and ISCO treatment (Figure 2A). Nutrient levels were overall low. PO_4_ and NOX were below the detection limit (0.04 μM for PO_4_ and 0.4 μM for NOX). Silicate levels dropped from t_0_ to t_2_ from maximum 7.20 μM to minimum 5.52 μM with no major differences between ISCO and HICO treatments (Figure 2B). Concentration of eCA in seawater was measured in quadruplicates from t_0_ to t_2_. Methods and data for this experiment were reported in the paper and supplement material of Mustaffa et al. (2017). Indeed, the concentration of eCA decreased faster in the HICO treatment compared to the ISCO treatment (Supplementary Figure 2 of Mustaffa et al. (2017)). ANOVA demonstrated significant differences for the eCA concentration (*F*=9.51, *df*=4, *p*=0.00049). Means of eCA concentrations were not significantly different between ISCO and HICO treatment at t_0_, but weakly significant at t_1_ (Tukey HSD, *p*=0.025). Missing data for the HICO treatment at t_2_ were removed for the statistics (and not assumed to be equal to zero).

**Figure 2:**
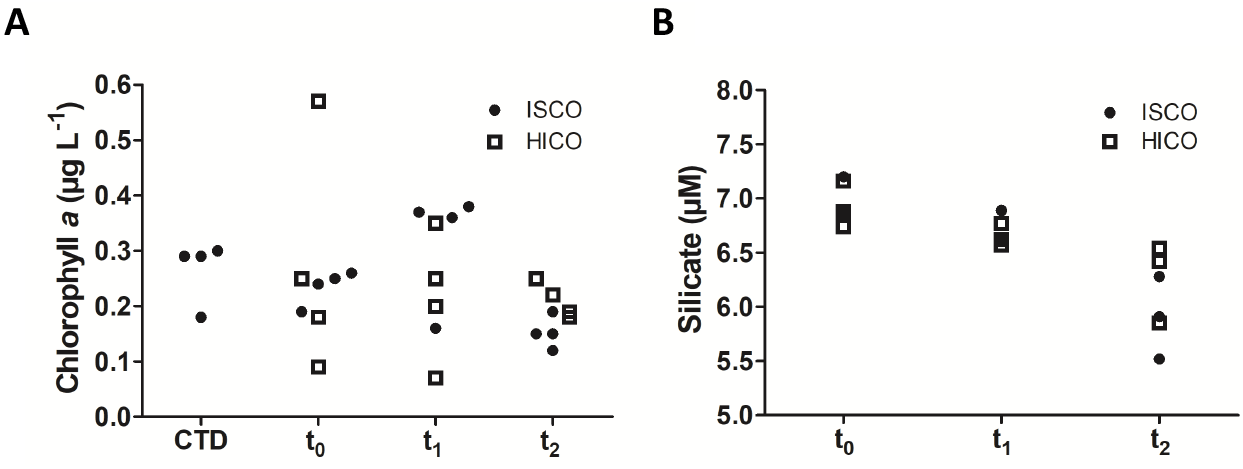
A) Chlorophyll *a* concentration and B) silicate (μM) measurements during the experiments (n=4).

### 3.3 Plankton alpha and beta diversity in response to acidification

The one-way ANOVA indicated weak significant differences between the mean species diversities of the different sample types (*F*=4.62, *df*=5, *p*=0.014). Species diversity analyses based on Shannon-Wiener index (Figure 3) showed that at t_0_ DNA-based OTUs showed a higher diversity measure in the ISCO (mean: 3.6 ± standard deviation: 0.98, n=3) compared to the HICO treatment (2.3 ± 0.53, n=3), which was not significant (Tukey HSD, *p*=0.1). Such differences were less pronounced for the cDNA-based OTUs, where both measures were equally high (ISCO: 4.5 ± 0.16, n=3; HICO: 4.4, n=2). For t_2_, the Shannon-Wiener index differences between the acidification treatments based on DNA-based OTUs were less strong compared to t_0_ (ISCO: 3.8 ± 0.46, n=3; HICO: 3.2 ± 0.44, n=3) and even more similar for cDNA-based OTUs (ISCO: 4.3 ± 0.05, n=3; HICO: 3.8 ± 0.53, n=3). It follows that OTUs based on DNA sequencing (reflecting abundant taxa) seem slightly more prone to acidification effects because they have a lower Shannon-Wiener index compared to those based on cDNA sequencing (reflecting active taxa), though all differences between treatment pairings were not significant based on Tukey HSD test.

**Figure 3:**
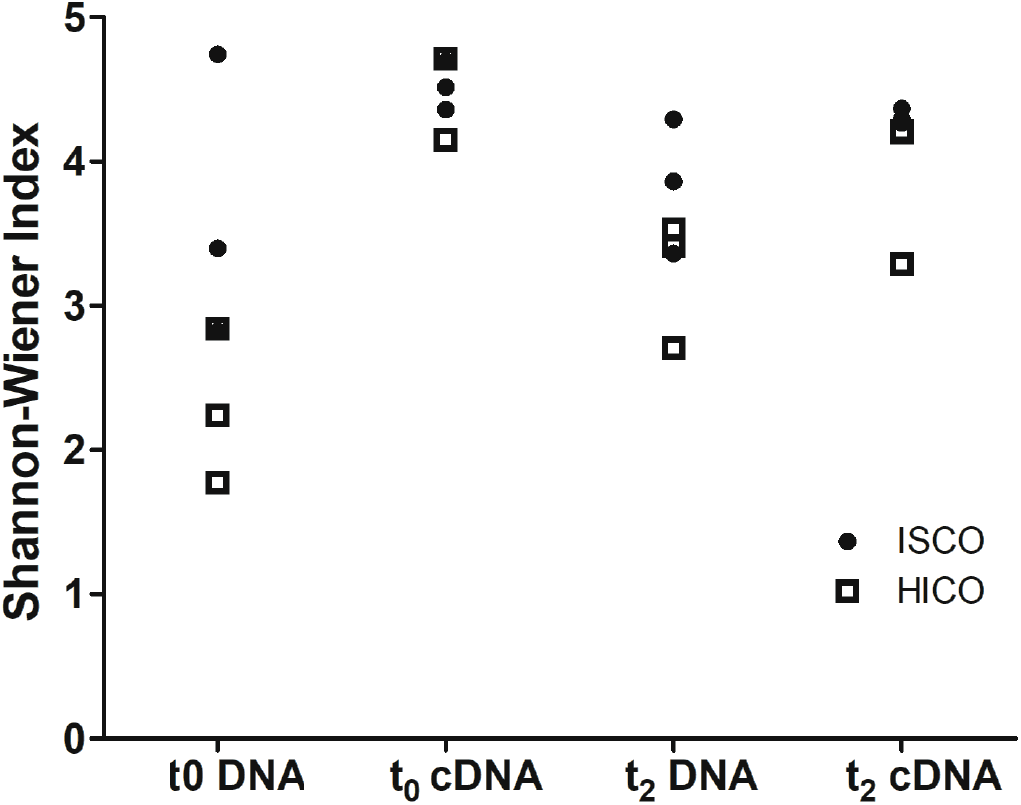
Calculated Shannon-Wiener indices (species richness) of ISCO and HICO-treated samples. HICO= high *p*CO_2_, ISCO= *in situ p*CO_2_

The community composition of each sample (Figure 4) revealed that the most striking differences in cDNA- and DNA-based OTUs between HICO and ISCO treatments at t_0_ were based on the different relative abundance of crustaceans. Overall, diversity of phyla appeared comparable between replicates (Figure 4). However, the community composition plots of the individual groups (diatoms, dinoflagellates etc.) showed profound differences in relative abundances between t_0_ and t_2_ (Supplementary Figures S2 – S12). Many OTUs that were undetectable at t_0_ reached considerable abundances at t_2_. For instance, after 48 hours, active and abundant OTUs assigned to uncultured kinetoplastids and especially the genus *Bodonidae* sp., the ciliate *Eutintinnus fraknoi*, the cercozoan *Massisteria marina* and the choanoflagellate *Calliacantha* sp., strongly increased in relative abundance from t_0_ to t_2_ (Supplementary Figures S4, S5, S6, S12). On the other hand, OTUs assigned to dinoflagellates and some haptophytes, e.g., *Phaeocystis antarctica* decreased in relative abundance, whereas other haptophytes, e.g., an OTU related to *Pavlova gyrans*, became more abundant during the incubation period (Supplementary Figures S3 and S7).

**Figure 4:**
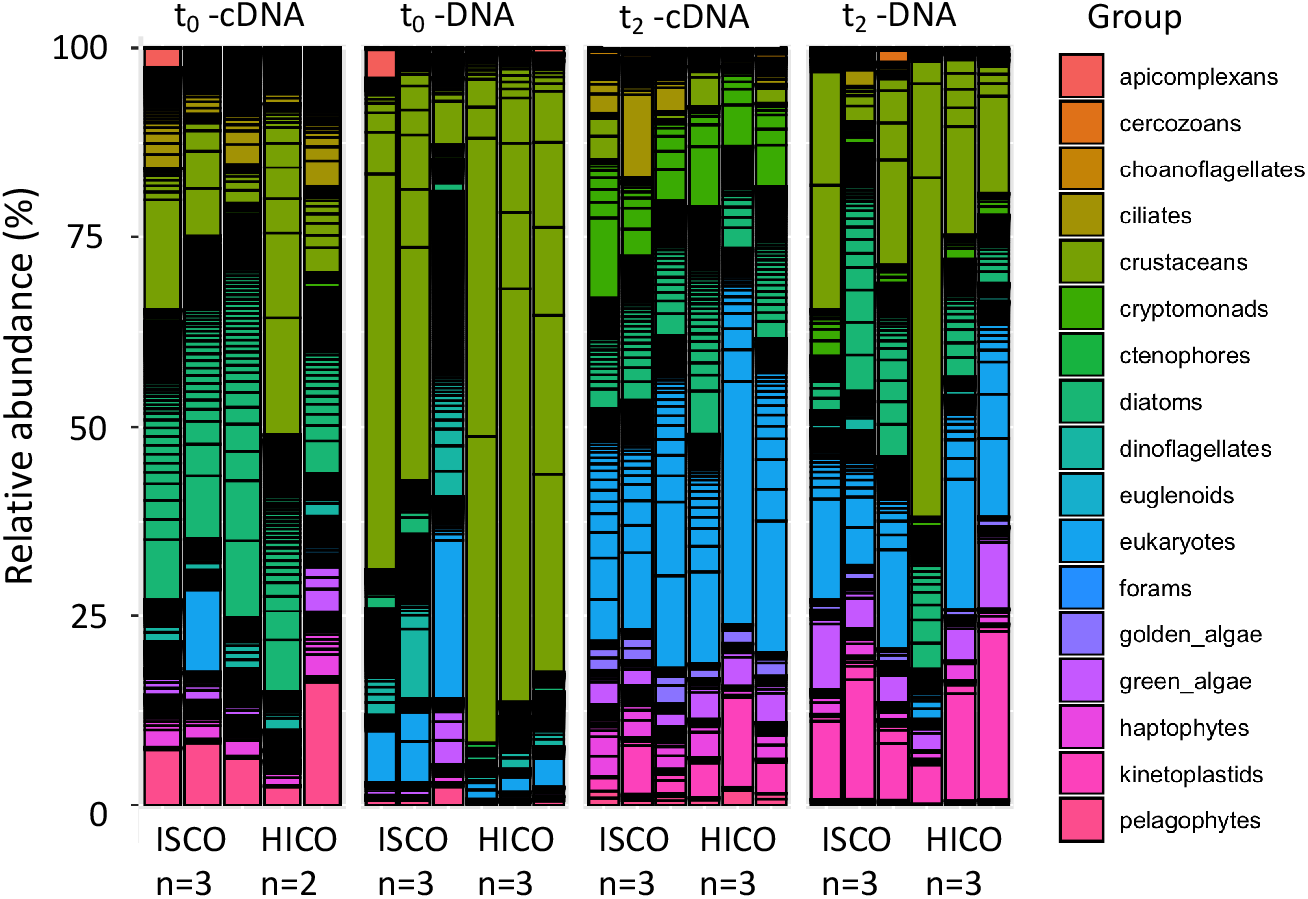
Community composition of taxa expressed as relative abundance (%) of operational taxonomic units (OTUs) for ISCO and HICO-treated samples of three replicates (with exception of one sample).

In agreement with these abundance changes of different taxa, the NMDS plot (Figure 5, stress= 0.070) indicated that four distinct community cluster were formed. The clusters could be mostly distinguished based on the nucleic acid type used as PCR template (DNA or cDNA) and the incubation period (t_0_ versus t_2_). Differences between HICO and ISCO treatment did not reveal strong effects at the phylum level, although at t_0_, the two treatments showed some separation among the DNA-based OTUs. Communities based on cDNA and DNA resembled each other more at t_2_ than at t_0_, and this observation was independent from the acidification treatment.

**Figure 5:**
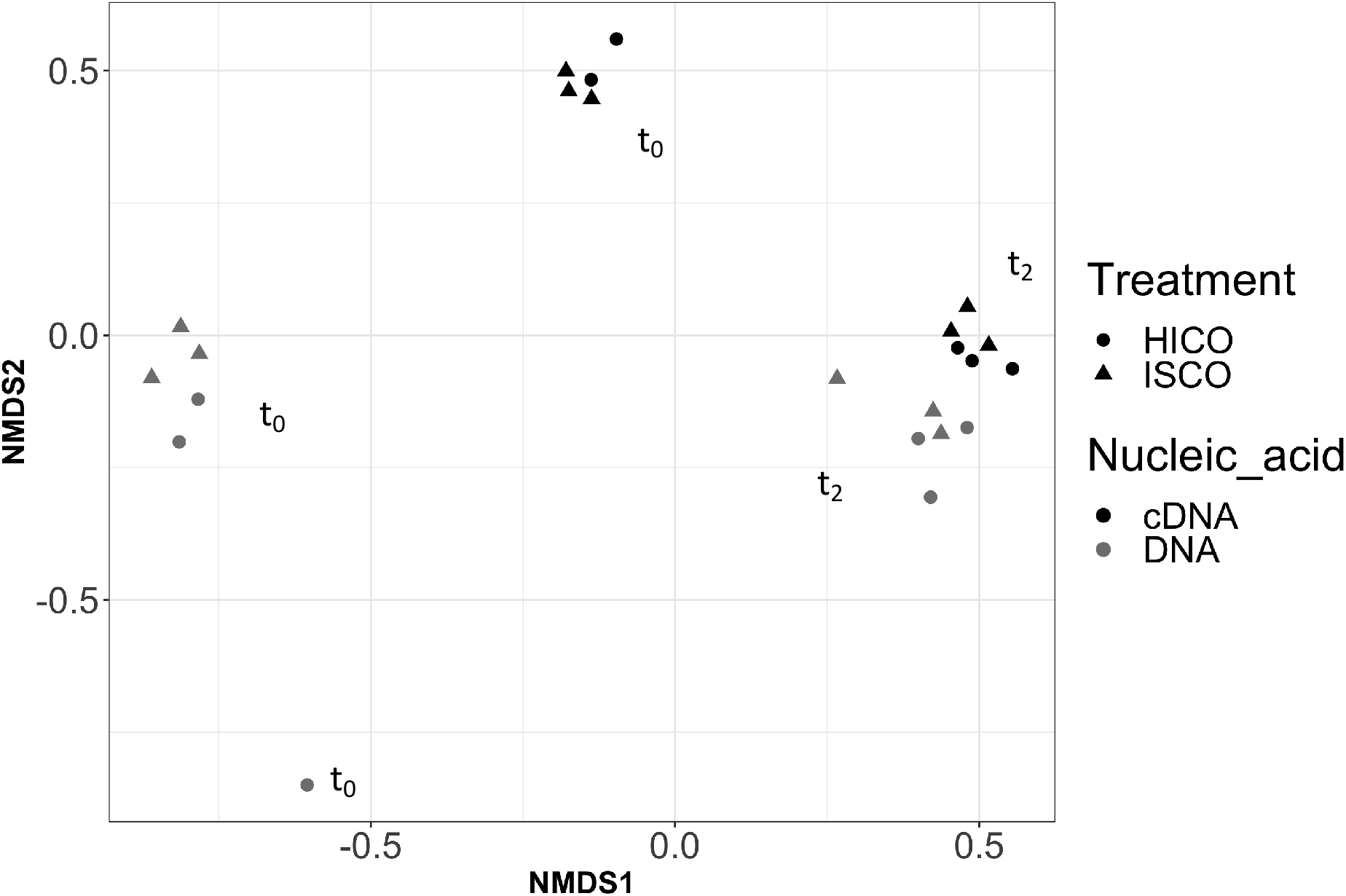
Non-metric multidimensional scaling (NMDS) plot calculated using Bray–Curtis dissimilarity. Circles indicate sample clusters. Stress=0.070, HICO= high *p*CO_2_, ISCO= *in situ p*CO_2_

### 3.4 OA effects on the OTU level

Despite the fact that the acidification treatment induced no major differences at the phylum level (Figure 4), significant differences (*p*≤0.05) were detectable at the OTU-level (Table 1). In most cases, the HICO treatment was associated with a lower relative abundance of significantly affected OTUs compared to the ISCO treatment.

**Table 1:**
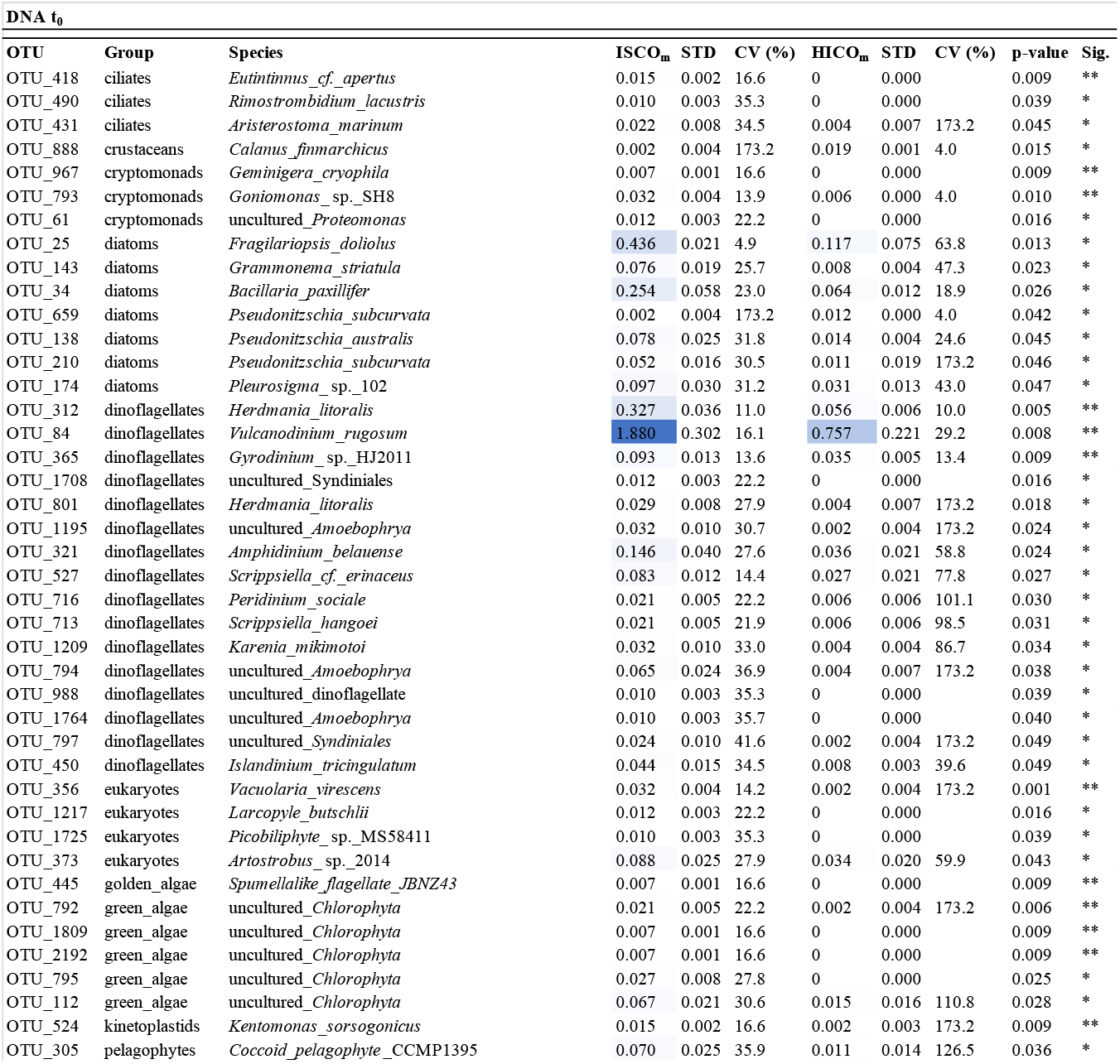
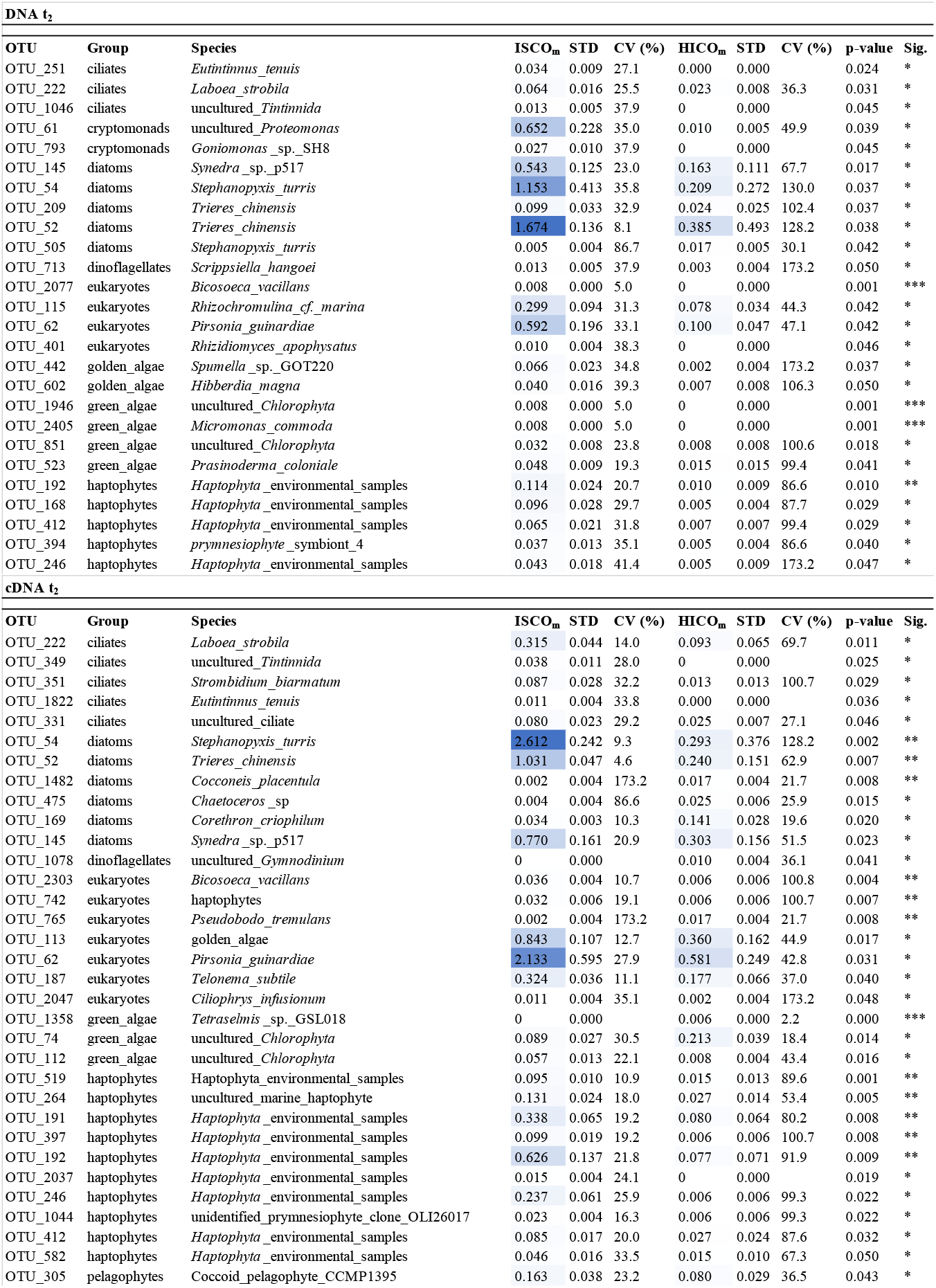
Operational taxonomic units (OTUs) that are significantly affected (p≤0.05) by the ocean acidification treatment (n=3 for each group). The value in the ISCOm and HICOm column represents the mean of relative abundance (%) for three biological replicates. Asteriks indicate significance (Sig.) level (*= p≤0.05, **= p≤0.01, ***= p≤0.001). CV=coefficient of variation, STD=standard deviation

For crustaceans, a higher relative abundance of the copepod species *Acartia longiremis* and *Calanus finmarchicus* was found among the DNA-based OTUs in the HICO treatment compared to the ISCO treatment (Supplementary Figure S8). For instance, the increase in relative abundance of an OTU assigned to *Calanus finmarchicus* was statistically significant in the HICO treatment at t_0_ (ISCO: DNA: mean=0.002 %, STD=±0.004 %, HICO: DNA: mean=0.02 % ±0.0007 %, *p*=0.015).

At t_2_, we sometimes found the same OTUs to be significantly affected by the OA treatment among the DNA and cDNA-based OTUs, e.g. for the diatoms *Trieres chinensis, Stephanopyxis turris* (OTU_54) or *Synedra* sp. (Table 1). All significantly affected taxa (based on Welch t-test) are shown in Table 1.

## 4. DISCUSSION

### 4.1 OA effects are species or even ecotype-specific

Increased atmospheric CO_2_ burden due to enhanced anthropogenic fossil fuel emissions and associated OA is one of the major global issues of our time. The effects of OA on natural eukaryotic plankton communities, especially from the tropical Timor Sea are poorly understood, and our data show that the HICO treatment influenced the relative abundance of certain abundant and active taxa from this oceanic region. Since the copy number of the small subunit rRNA genes is known to correlate with the biovolume of marine diatoms and can reach considerable numbers per cell in this group (Godhe et al. 2008) and also in ciliates (Gong et al. 2013), we are aware about the difficulties of using this gene for the interpretation of single-celled protist numbers. However, it is impossible to know the exact copy number of the 18S rRNA (gene) for each OTU belonging to different protists of a mixed community of which many species are uncultured or even unknown. We made comparisons on the relative abundance of an OTU by comparing two treatments, meaning that even if an OTU species-specifically contains multiple 18S rRNA gene copies, it will probably do so in both treatments and is thus comparable.

While significant CO_2_ effects were not immediately apparent at higher taxonomic levels (Figure 4), they became visible at the OTU level. We expected that phytoplankton OTUs would be most sensitive to changes in CO_2_ due to their inorganic carbon requirements for primary production and found responses to OA in diatoms to be highly species- or even ecotype specific. Many diatoms affected by the HICO treatment decreased in relative abundance as a response including *Trieres chinensis, Synedra* sp., *Fragilariopsis doliolus, Pseudonitzschia australis, Pseudo-nitzschia subcurvata* (OTU_210), *Stephanopyxis turris* (OTU_54), and *Pleurosigma* sp. The diatom *Stephanopyxis turris* (OTU_54) showed a decrease in relative abundance in response to the HICO manipulation both in the DNA-and the cDNA-based investigation after 48 hours, most likely indicating that abundance and activity were both reduced by the treatment. This further confirms that the significant effect was real and not a false-positive random finding. A few diatoms also increased in abundance in the HICO treatment such as *Cocconeis placentula, Chaetoceros* sp., *Corethron criophilum, Grammonema striatula*, another *Pseudo-nitzschia subcurvata* (OTU_659) and another *Stephanopyxis turris* (OTU_505), matching the generally constant chl *a* concentration. Our results are in agreement with Schulz et al. (2017) postulating that CO_2_ effects on diatoms are more likely to become visible on the species rather than on the phylum- or other higher taxonomic level.

Many studies of OA effects on natural microbial communities focus on polar waters complicating direct comparison to our findings from a tropical region. By investigating phytoplankton populations from the Ross Sea, Southern Ocean, Tortell et al. (2008) reported that growth of larger chain-forming diatoms such as *Chaetoceros* spp. increased in abundance in response to high CO_2_ (800 μatm), while the small pennate diatom *Pseudo-nitzschia subcurvata* decreased in this treatment. This finding matches our results apart from the fact that we had two different OTUs assigned to *Pseudo-nitzschia subcurvata*, and both had contrasting responses to the treatment at t_0_ and that our HICO treatment was much higher (*p*CO_2_=1823 μatm). Also, we cannot conclude from our results that different morphologies of diatoms (centric vs. pennate) have different advantages in facing high and low CO_2_ environments as proposed by Tortell et al. (2008), because in our study OTUs related to both morphology types decreased in relative abundance.

We also found contrasting responses to OA by two OTUs being both assigned to *Stephanopyxis turris*. This leads to the conclusion that very closely related diatom species (97% sequence identity) can have very different responses to OA. Different responses of the same model species *Emiliania huxleyi* (but from different ecotypes) to carbonate chemistry manipulations were previously described (Ridgwell et al. 2009). An ecotype refers to a genetically distinct geographic population within a species. Ecotype-specific OA responses might lead to conflicting results between studies and explain the contrasting responses we found for OTUs assigned to *Stephanopyxis turris* and *Pseudo-nitzschia subcurvata*.

Immediate responses to OA were also observed for the dinoflagellate *Vulcanodinium rugosum*, which decreased in relative abundance at t_0_. *V*. *rugosum* is known for being a harmful algae and producing pinnatoxins causing shellfish poisoning (Rhodes et al. 2011). While it is known that *V. rugosum* is thermophile and euryhaline (Abadie et al. 2016, Abadie et al. 2018), nothing is known about its response to to OA. The immediate response and decrease in relative abundance among DNA-based OTUs from mean ± STD=ISCO DNA mean: 1.9% ± 0.30 to HICO DNA mean: 0.76% ± 0.22 (n=3, Welch t-test, *p*=0.0083) suggests that this dinoflagellate was sensitive to the HICO treatment. Effects of CO_2_-induced OA on dinoflagellates that are able to form (harmful) algal blooms have been previously studied, again showing very species-specific responses. For instance, biomass production in calcareous *Scrippsiella trochoidea* decreased, while toxic *Alexandrium tamarense* did not respond to an OA treatment (Eberlein et al. 2014).

Two copepod OTUs were BLAST-assigned to *Acartia longiremis* and *Calanus finmarchicus*. Although both species have their respective main distribution in the North Atlantic, we decided to keep these names here because there are some records of these species in the southern hemisphere in OBIS (https://obis.org). Both copepods showed sudden increases in relative abundance in response to the HICO treatment at t_0_. The increases were striking but mostly not significant, most likely due to the variation between replicates. We can only speculate about the reason for the immediate response. It has been shown that lowered seawater pH can influence copepod reproduction, hatching and development in some species (Kurihara et al. 2004, Mayor et al. 2007) whereas it had no significant effect on physiological parameters or grazing of copepods in other studies (Kurihara & Ishimatsu 2008, Mayor et al. 2012, Isari et al. 2015, Hildebrandt et al. 2016). This implies that OA effects are very species and context-specific and also dependent on the geographic origin of the copepod population (Thor & Oliva 2015), the life stage (Cripps et al. 2014), gender (Cripps et al. 2016), and even food quality provided (McLaskey et al. 2019). Samples that were most affected by the HICO treatment at t_0_ could have randomly contained a particular CO_2_-vulnerable life stage or gender of copepods. Or copepod carcasses and fecal pellets present in the samples were more slowly degraded in the HICO compared to the ISCO treatment.

By assigning affected taxa to different OTU abundance ranges (Table 2), we found that the proportion of HICO-treatment affected OTUs was lower compared to the ISCO treatment, especially in the high abundance ranges. This suggests that the HICO treatment overall had a suppressive effect on the relative abundance of taxa. For instance, OTUs that reached an minimum abundance of 9.4% in the ISCO treatment in the 1-100% abundance range, were undetectable at such a high abundance in the HICO treatment. Not surprisingly, more OTUs were found in the lowest abundance range in the HICO treatment, which might be attributable to the pH manipulation.

**Table 2:** Proportion (%) of significantly CO_2_-affected OTUs within the given relative abundance range for each sample

| Rel. Abundance<br>range of OTUs | ISCO-DNA-t0 | HICO-DNA-t0 | ISCO-DNA-t2 | HICO-DNA-t2 | ISCO-cDNA-t2 | HICO-cDNA-t2 |
| --- | --- | --- | --- | --- | --- | --- |
| 1-100% | 9.4 | 0.0 | 12.0 | 0.0 | 13.4 | 0.0 |
| 0.1-1% | 6.1 | 8.3 | 7.6 | 8.1 | 9.5 | 9.6 |
| 0.01-0.1% | 6.4 | 5.1 | 6.9 | 3.3 | 7.8 | 6.8 |
| 0.001-0.01% | 3.2 | 6.3 | 2.3 | 6.4 | 2.5 | 7.9 |
| 0-0.001% | 0.0 | 0.9 | 0.0 | 0.5 | 0.1 | 0.2 |

### 4.2 Adaptations to the ocean acidification treatment

The NMDS plot revealed that eukaryotic plankton community differences between DNA and cDNA-based OTUs became smaller at t_2_ compared to t_0_ (Figure 5). The incubation conditions (heat, low oxygen, and grazing pressure) likely selected for specific taxa that could resist those conditions and became abundant and active, while it led to a decreased abundance or even extinction of others. The “bottle enclosure effect” exerted by incubations in closed systems (Gieskes et al. 1979) includes decreasing biomass of picophytoplankton, increasing biomass of heterotrophic bacteria (Calvo-Diaz et al. 2011), and increases the likelihood for host cell-virus encounters with potential to alter the community composition (Haro-Moreno et al. 2019). However, containment of natural seawater for 48 hours led to only minor changes in species richness (Figure 3, Shannon-Wiener Index) as previously reported (Countway et al. 2005).

We found an increase in the relative abundance of certain taxa from t_0_ to t_2_, mainly of kinetoplastids, cercozoans, cryptomoands and choanoflagellates (Supplementary Figures S5, S6, S9, S12), while other taxa such as dinoflagellates tended to decrease in relative abundance. The decrease in relative abundance of dinoflagellates might be due to increased grazing or to enhanced vulnerability towards incubation conditions and associated community changes.

The fact that abundant and active eukaryotic communities became more similar with increased incubation time emphasizes that several members of the planktonic community were strongly affected by the incubation conditions. Since the effect occurred after 48 hours (t_2_) and independent of the OA treatment, our data show that longer bottle incubations to test for OA effects have to be conducted with great caution. If a reliable presentation of the *in situ* community is needed, direct processing at t_0_ is required as has been recently suggested for prokaryotic communities (Haro-Moreno et al. 2019). On the other hand, the “bottle enclosure effect” can also be systematically used to allow rare and underrepresented populations to flourish. OA studies often apply acclimation periods with gradually increasing CO_2_ to let organisms slowly adjust to new pH conditions (Hancock et al. 2018), which is probably a more realistic representation of OA than sudden pH manipulations. However, here we used acute elevations of seawater *p*CO_2_ to obtain a better understanding of the short-term sensitivity of planktonic organisms and because rapid (within a few days) shifts in picoeukaryote communities due to OA were previously detected (Meakin & Wyman 2011).

Adaptation of the planktonic microbial eukaryotes to the OA treatment also became visible at the enzymatic level. The down-regulation of carbon concentrating mechanisms processes in response to high CO_2_ serves as an energy saving mechanism, which has been previously observed in model phytoplankton species (Wu et al. 2010, Moroney et al. 2011) and natural phytoplankton communities from Antarctica (Young et al. 2015, Deppeler et al. 2018). Carbonic anhydrases, zinc-containing metallo-enzymes that catalyze the hydration of CO_2_ into bicarbonate, are important components for the functioning of carbon concentrating mechanisms under CO_2_-limiting conditions (Mondal et al. 2016). Likewise, our data show a rapid decline in the concentration of extracellular carbonic anhydrase (eCA) in response to the HICO treatment in the (autotrophic) plankton community indicating an adaptation to the manipulation. The eCA concentration dropped more slowly in the ISCO compared to the HICO treatment (Supplementary Fig.2 of Mustaffa et al. (2017)). Dropping of eCA concentration in the ISCO treatment probably happened because of a depletion in phytoplankton biomass due to grazing in closed bottles and associated accumulation of respired CO_2_ suppressing carbon concentrating mechanisms.

Despite constant pumping of fresh ocean water into the sampling pool, heating of incubation bottles during the day because of the tropical climate (Supplementary Figure S1) was a clear limitation of our experiment. Previous studies did not detect a synergetic effect of OA and warming on plankton communities (Paul et al. 2015, Horn et al. 2016). But to exclude additional effects by warming, e.g., the decrease of heterotrophic flagellate biomass (Moustaka-Gouni et al. 2016), we decided to compare effects of OA treatment only within but not between t_0_ and t_2_. However, we assume that planktonic organisms were probably not adversely affected, i.e. killed, by the temperature at least until t_2_ because of the positive development of some groups and ongoing transcription of rRNA indicating active cell growth and activity (Schaechter et al. 1958, Poulsen et al. 1993, Lanzén et al. 2011).

OA effects on natural planktonic communities especially from tropical regions are poorly understood and require further investigation. Incubations over several days provide the advantage that they sometimes allow rare taxa to flourish (even if artificially), hence allowing effects of OA to be discovered that would have stayed obscure otherwise. OA might particularly affect the rare and some thermophilic taxa believed to cope well with climate change without considering their actual response to OA. Although being rare, some taxa could be of human interest, because they produce important metabolites or are somehow involved in the complex interactions of the oceanic food web. Diatoms and dinoflagellates as important primary producers and base of the food web seemed particularly vulnerable to OA and incubation-induced community changes. Overall, unravelling the interactive or opposed effects of a warming and simultaneously acidifying ocean on different eukaryotic species from natural planktonic communities is a difficult task due to differential intra- and interspecies responses. The consequences for whole trophic networks of a future ocean will consequently be even harder to predict.

## 5. ACKNOWLEDGEMENTS

We thank the captain and crew members of the *R/V* Falkor (cruise FK161010). We further acknowledge the assistance of Lea Oeljeschläger and Nur Ili Hamizah Mustaffa during sample processing. We further thank Claudia Wylezich for sharing her expertise and providing valuable feedback on our manuscript. This work was funded by the European Research Council (ERC) project PASSME (grant number GA336408). This is publication number 69 from the Senckenberg am Meer Metabarcoding and Molecular Laboratory.

## Supplementary Material

**Supplementary Figure S1:**
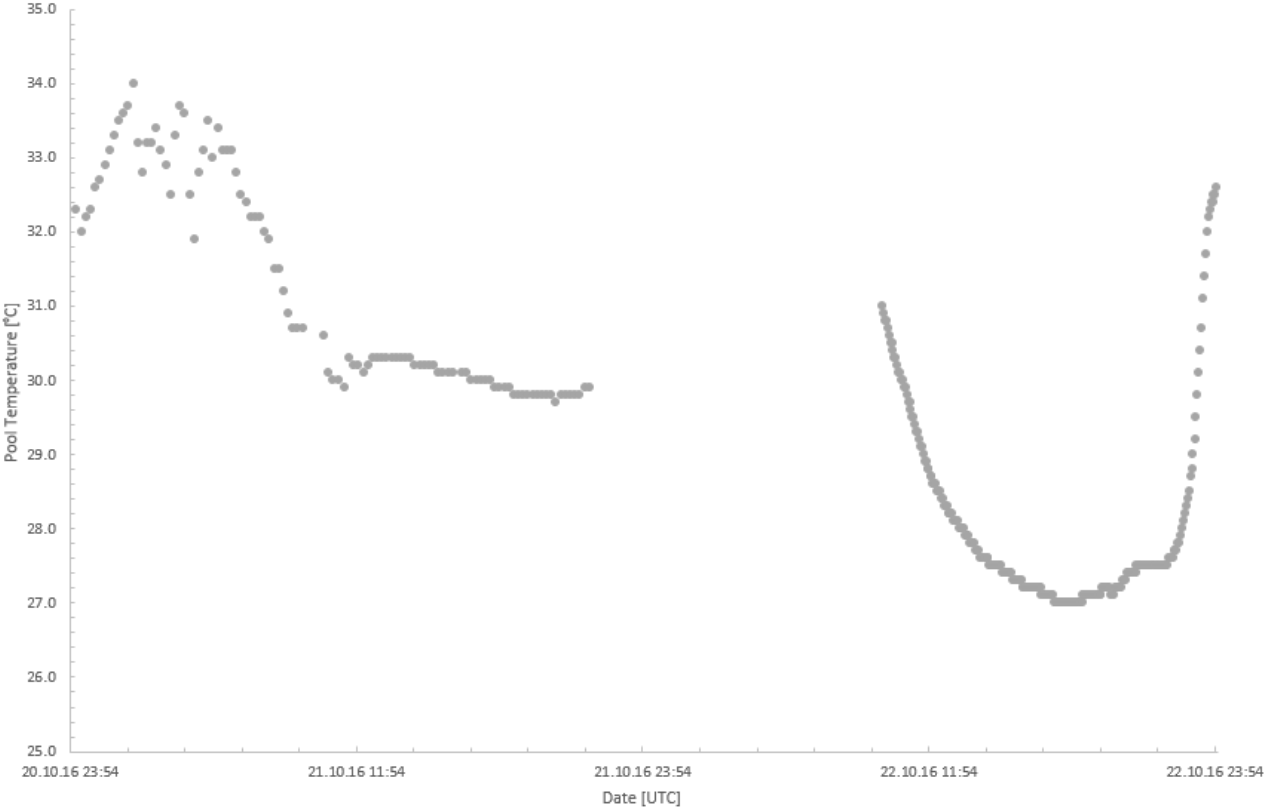
Temperature of pool water (°C) surrounding incubated bottles. Dates refer to t_0_ to t_2_, respectively. The data gap happened due to an unrecognized full memory of the measuring device.

**Supplementary Table S1:**
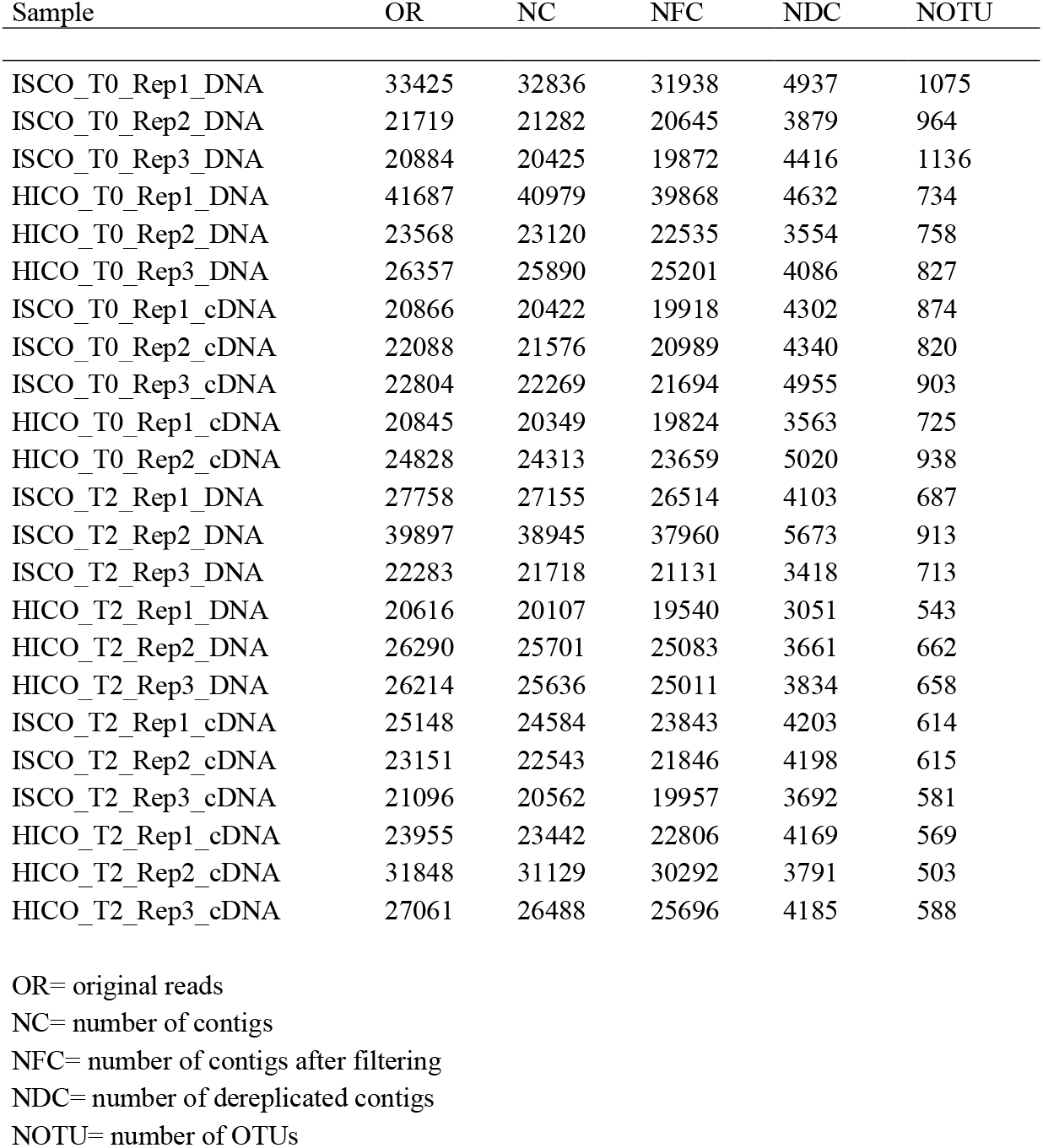
Sequencing statistic

**Supplementary Figure S2:**
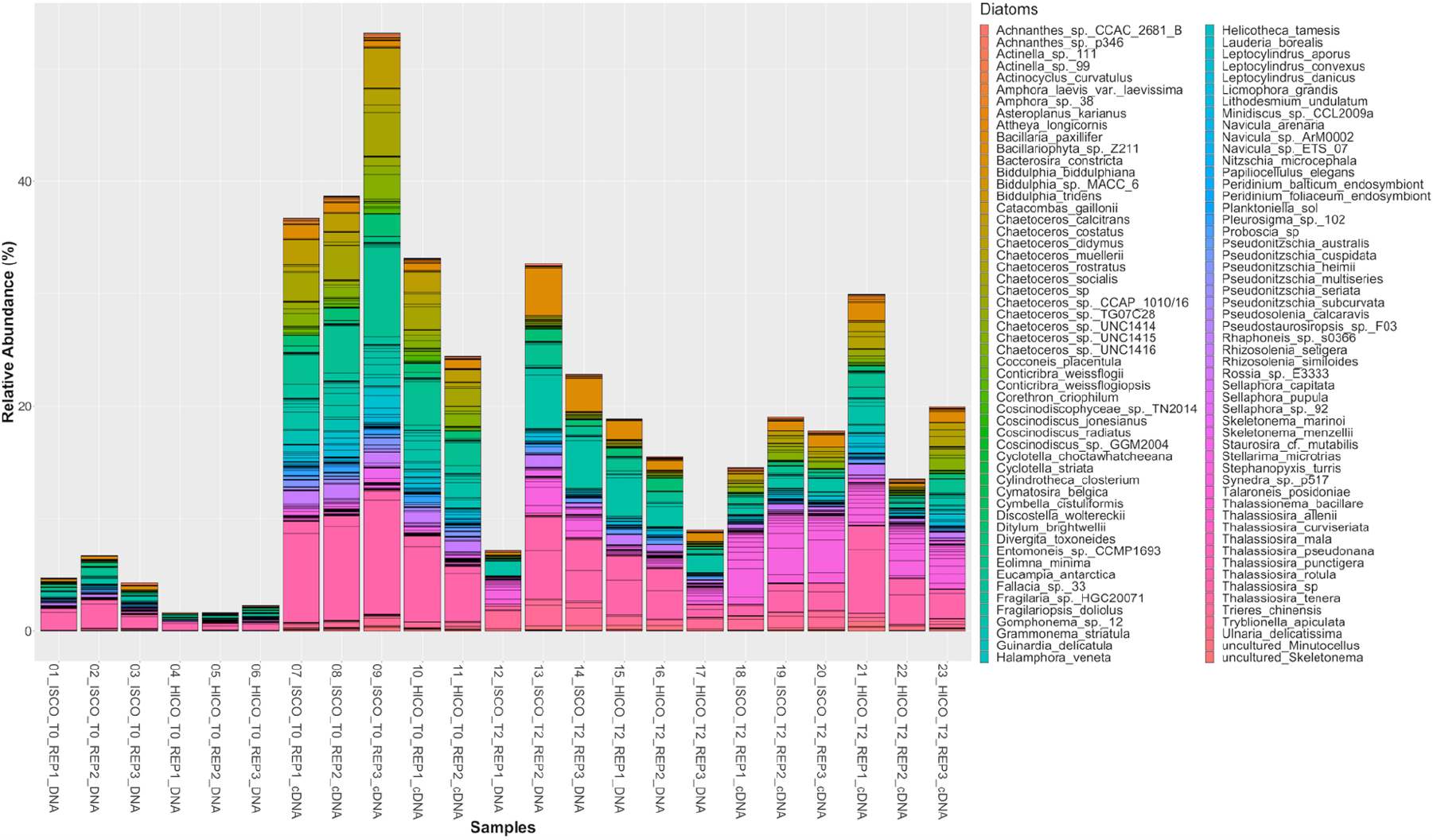
Relative abundance (%) of diatoms in ISCO (*in situ* CO_2_) and HICO (high CO_2_) treatment after immediate (t_0_) and 48 hours (t_2_) treatment exposure.

**Supplementary Figure S3:**
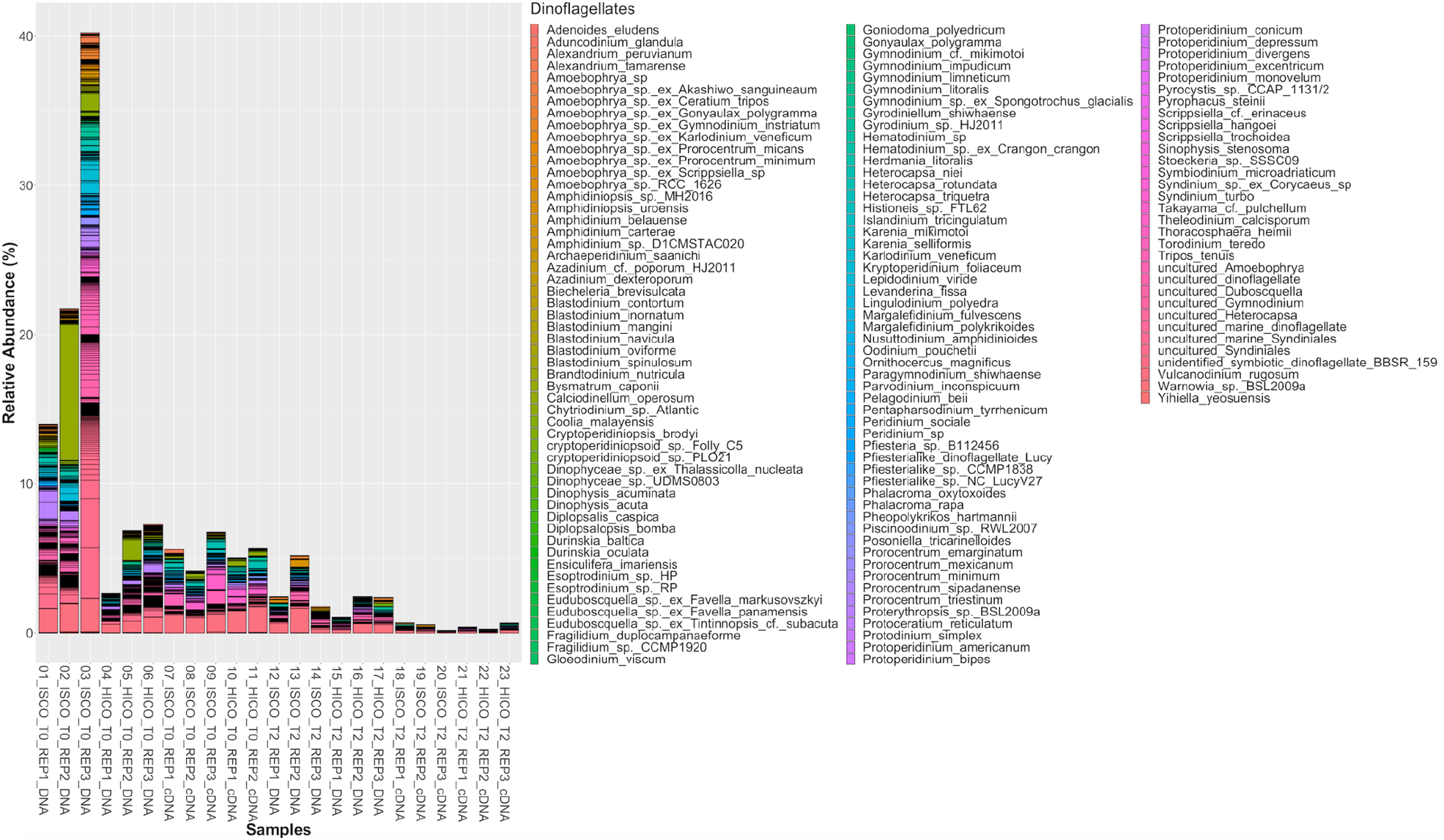
Relative abundance (%) of dinoflagellates in ISCO (*in situ* CO_2_) and HICO (high CO_2_) treatment after immediate (t_0_) and 48 hours (t_2_) treatment exposure.

**Supplementary Figure S4:**
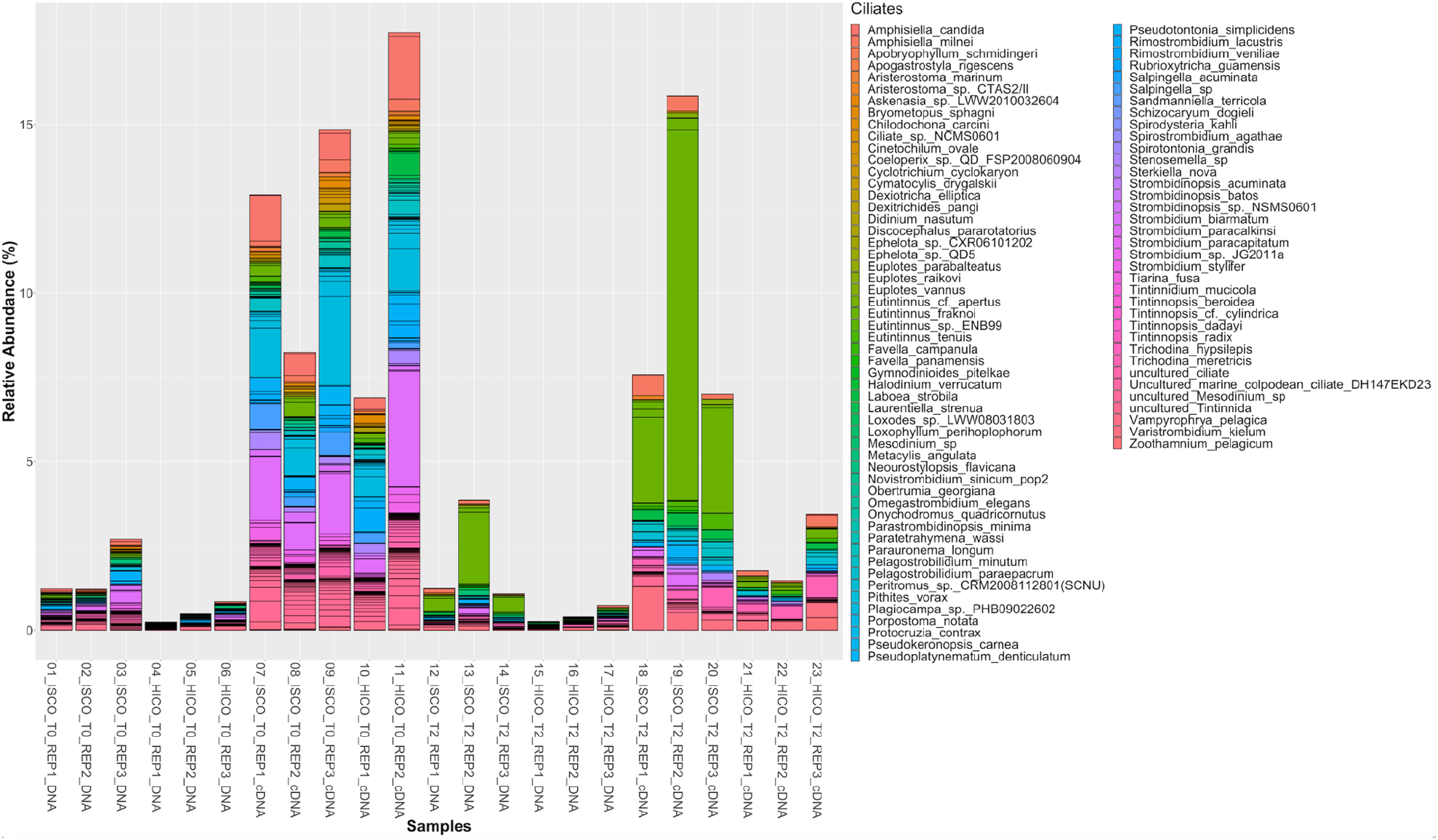
Relative abundance (%) of ciliates in ISCO (*in situ* CO_2_) and HICO (high CO_2_) treatment after immediate (t_0_) and 48 hours (t_2_) treatment exposure.

**Supplementary Figure S5:**
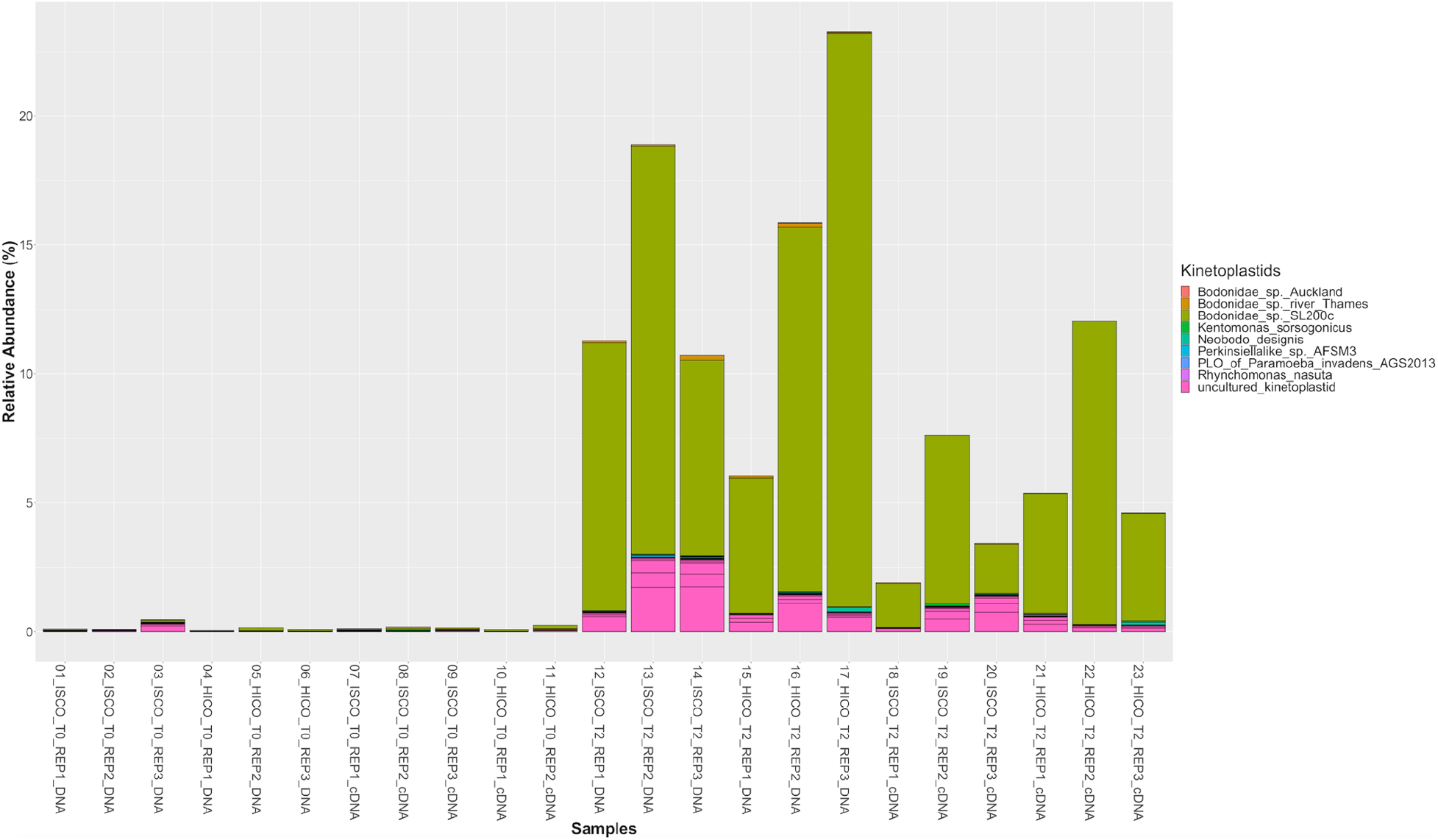
Relative abundance (%) of kinetoplastids in ISCO (*in situ* CO_2_) and HICO (high CO_2_) treatment after immediate (t_0_) and 48 hours (t_2_) treatment exposure.

**Supplementary Figure S6:**
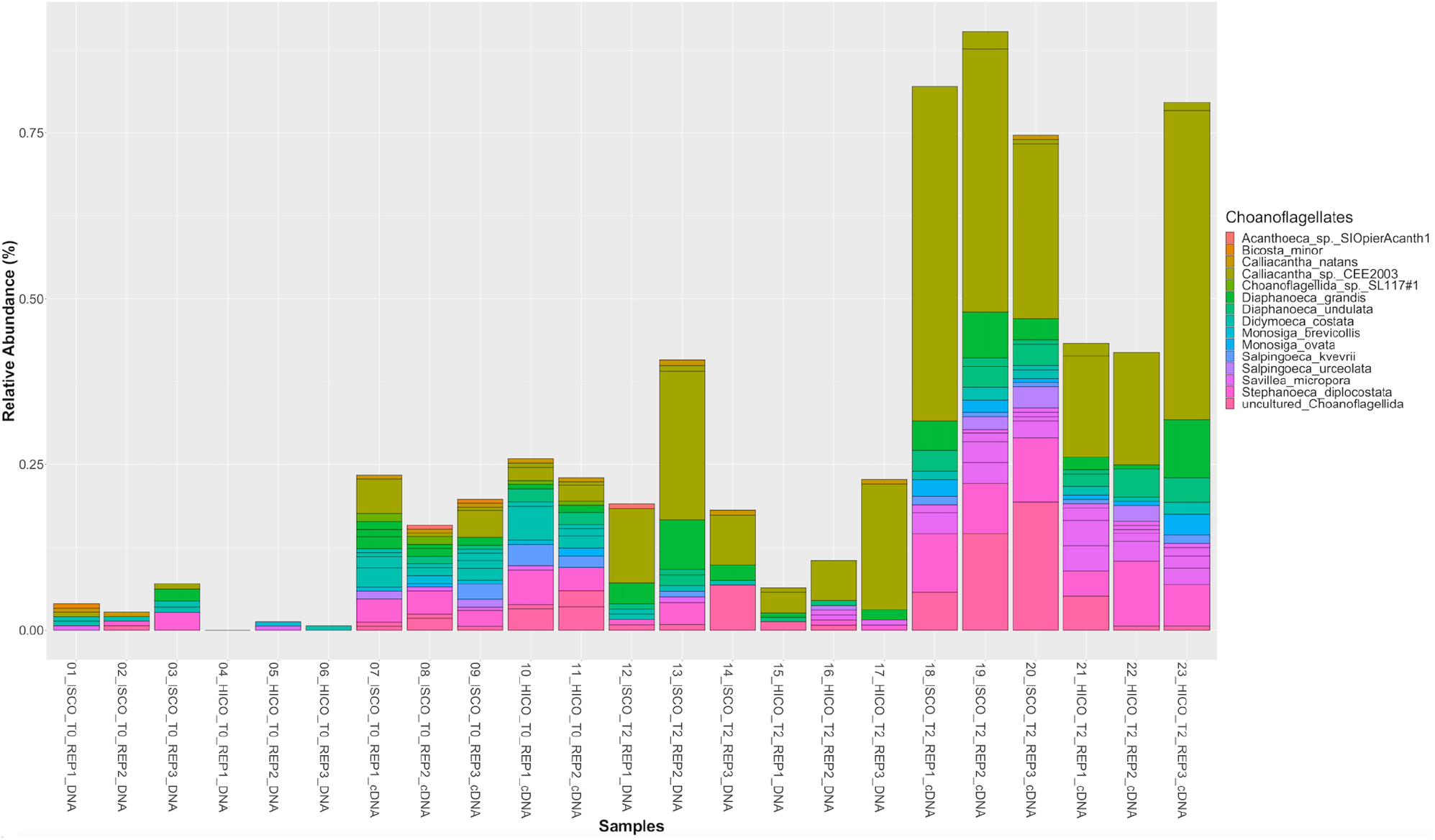
Relative abundance (%) of choanoflagellates in ISCO (*in situ* CO_2_) and HICO (high CO_2_) treatment after immediate (t_0_) and 48 hours (t_2_) treatment exposure.

**Supplementary Figure S7:**
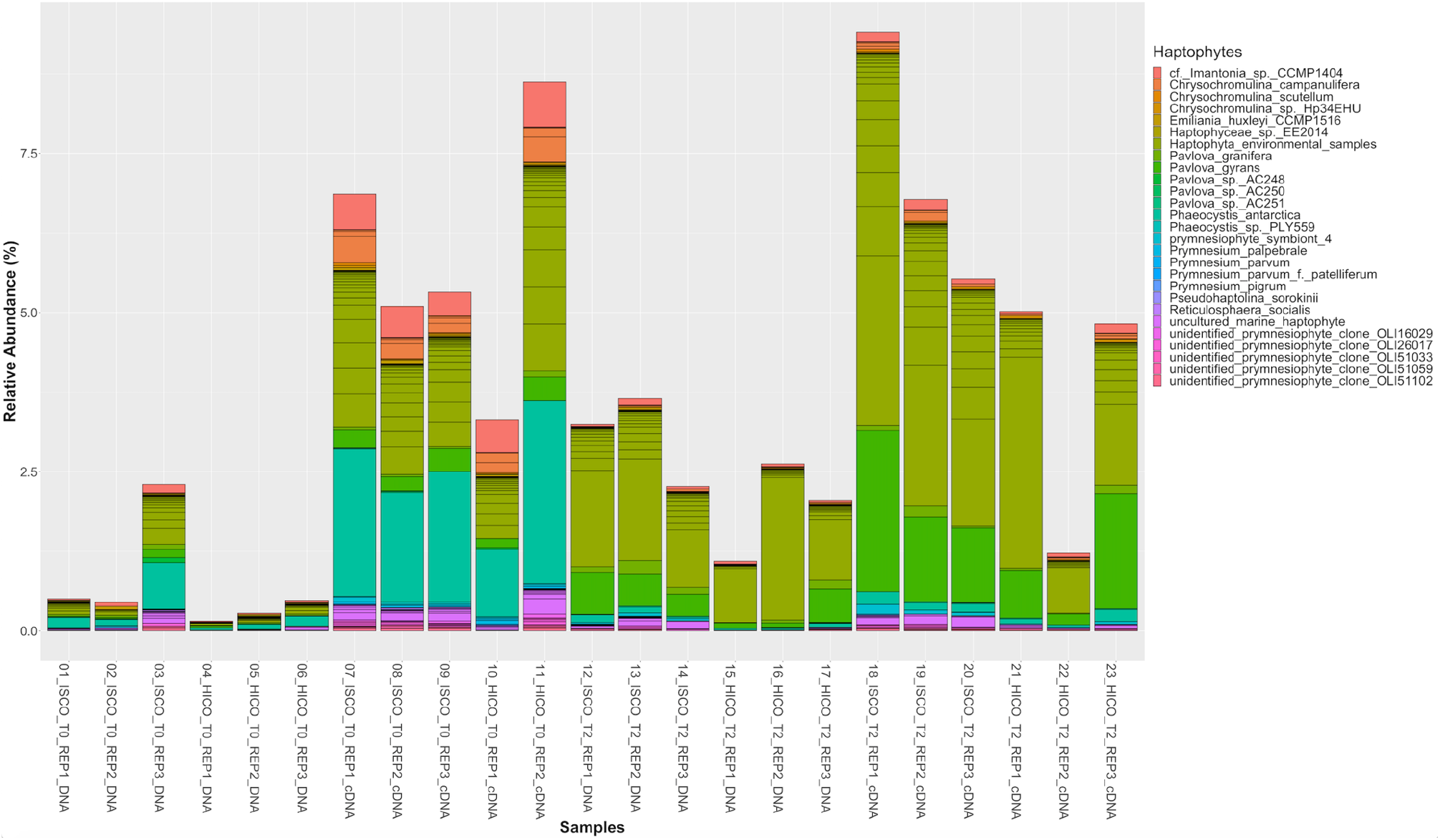
Relative abundance (%) of haptophytes in ISCO (*in situ* CO_2_) and HICO (high CO_2_) treatment after immediate (t_0_) and 48 hours (t_2_) treatment exposure.

**Supplementary Figure S8:**
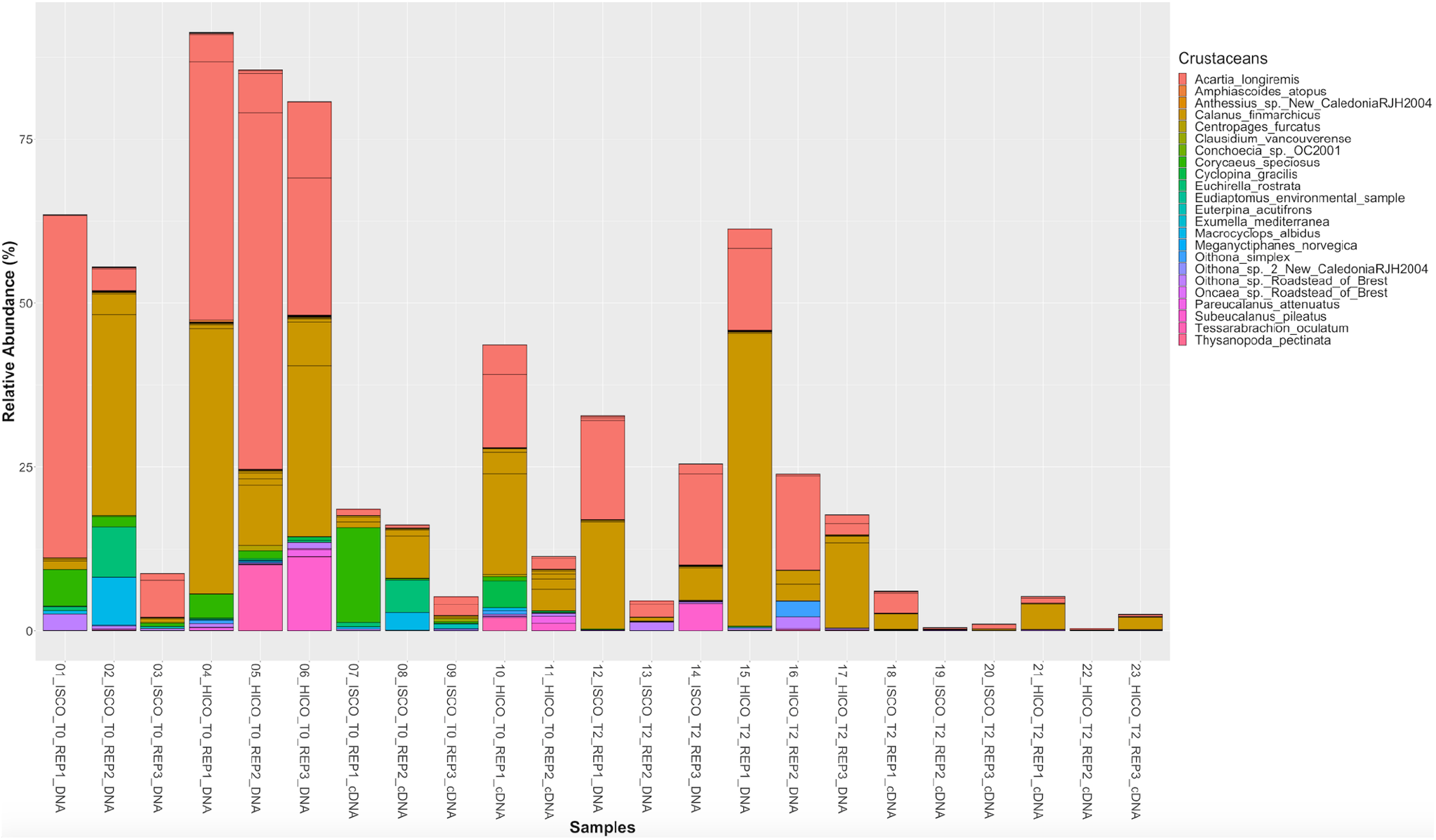
Relative abundance (%) of crustaceans in ISCO (*in situ* CO_2_) and HICO (high CO_2_) treatment after immediate (t_0_) and 48 hours (t_2_) treatment exposure.

**Supplementary Figure S9:**
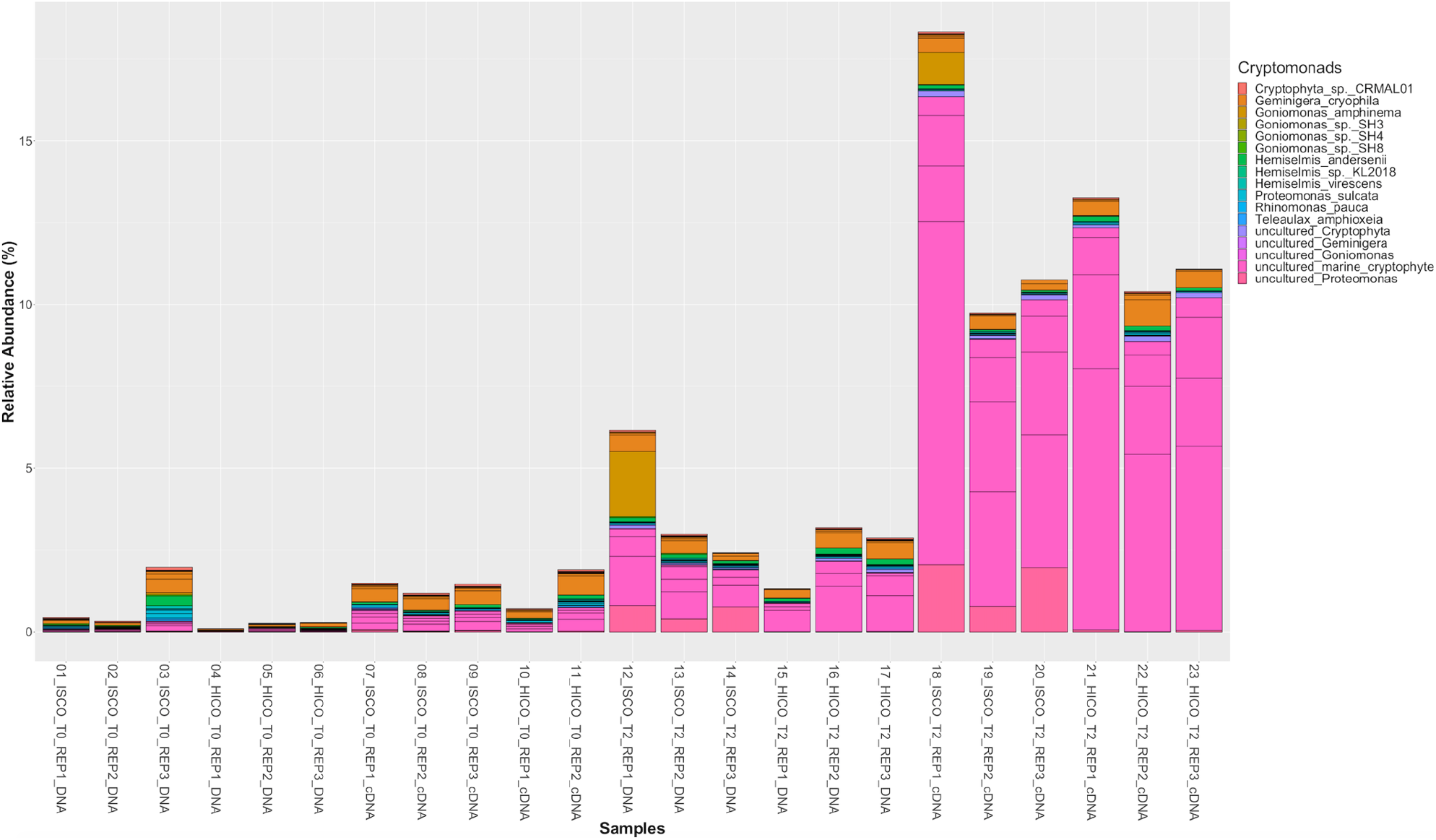
Relative abundance (%) of cryptomonads in ISCO (*in situ* CO_2_) and HICO (high CO_2_) treatment after immediate (t_0_) and 48 hours (t_2_) treatment exposure.

**Supplementary Figure S10:**
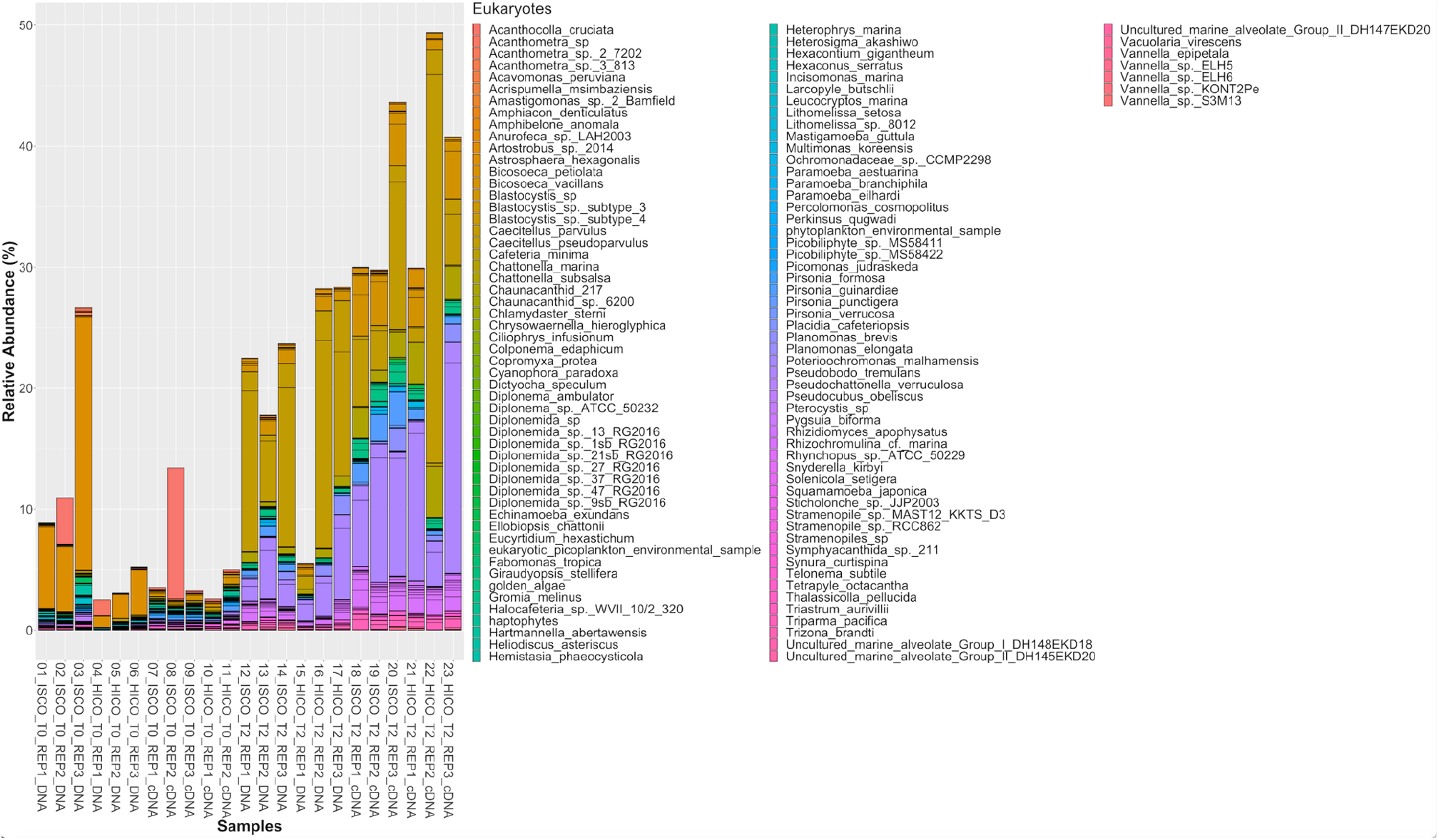
Relative abundance (%) of eukaryotes in ISCO (*in situ* CO_2_) and HICO (high CO_2_) treatment after immediate (t_0_) and 48 hours (t_2_) treatment exposure.

**Supplementary Figure S11:**
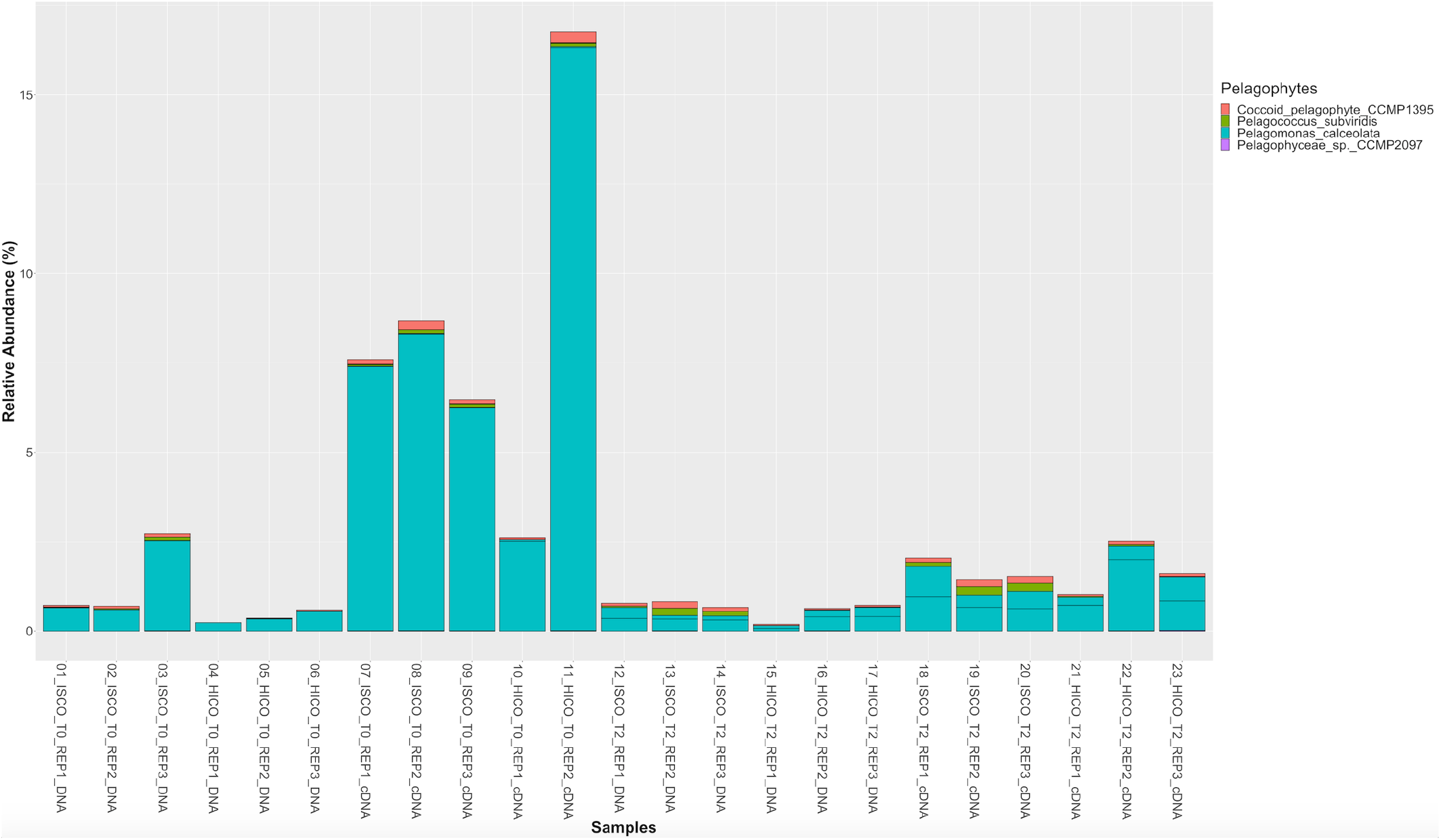
Relative abundance (%) of pelagophytes in ISCO (*in situ* CO_2_) and HICO (high CO_2_) treatment after immediate (t_0_) and 48 hours (t_2_) treatment exposure.

**Supplementary Figure S12:**
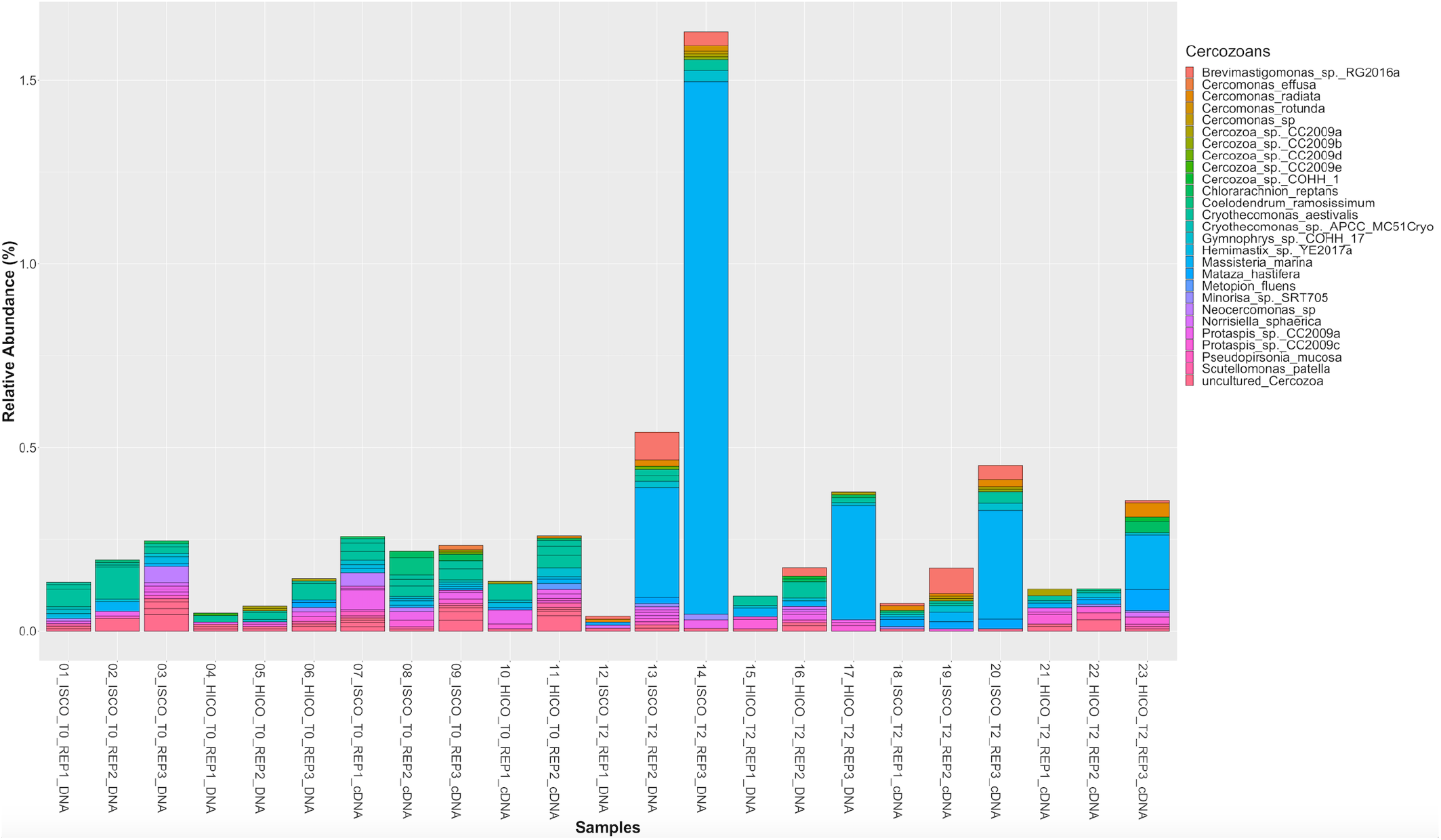
Relative abundance (%) of cercozoans in ISCO (*in situ* CO_2_) and HICO (high CO_2_) treatment after immediate (t_0_) and 48 hours (t_2_) treatment exposure.

